# Nicheformer: a foundation model for single-cell and spatial omics

**DOI:** 10.1101/2024.04.15.589472

**Authors:** Anna C. Schaar, Alejandro Tejada-Lapuerta, Giovanni Palla, Robert Gutgesell, Lennard Halle, Mariia Minaeva, Larsen Vornholz, Leander Dony, Francesca Drummer, Mojtaba Bahrami, Fabian J. Theis

## Abstract

Tissue makeup relies fundamentally on the cellular microenvironment. Spatial single-cell genomics allows probing the underlying cellular interactions in an unbiased, scalable fashion. To learn a unified cell representation that accounts for local dependencies in the cellular microenvironment, we propose Nicheformer, a transformer-based foundation model that combines human and mouse dissociated single-cell and targeted spatial transcriptomics data. Pretrained on over 57 million dissociated and 53 million spatially resolved cells across 73 tissues on cellular reconstruction, the model is fine-tuned on spatial tasks for spatial omics data to decode spatially resolved cellular information. Nicheformer excels in linear-probing and fine-tuning scenarios for a novel set of downstream tasks, in particular spatial composition prediction and spatial label prediction. We further show that existing foundation models trained on dissociated single-cell data alone are not capable of recapitulating the spatial complexity of cells in their microenvironments, indicating that multiscale models are required to understand complex local dependencies at scale. Nicheformer enables the prediction of the spatial context of dissociated cells, allowing the transfer of rich spatial information to scRNA-seq datasets. Overall, Nicheformer sets the stage for the next generation of machine-learning models in spatial single-cell analysis.

**Extended Abstract:** Tissue makeup and the corresponding orchestration of vital biological activities, ranging from development and differentiation to immune response and regeneration, rely fundamentally on the cellular microenvironment and the interactions between cells. Spatial single-cell genomics allows probing such interactions in an unbiased and, increasingly, scalable fashion. To learn a unified cell representation that accounts for local dependencies in the cellular microenvironment and the underlying cell interactions, we propose to generalize recent foundation modeling approaches for disassociated single-cell transcriptomics to the spatial omics setting.

Our model, Nicheformer, is a transformer-based foundation model that combines human and mouse dissociated single-cell and targeted spatial transcriptomics data to learn a cellular representation useful for a large variety of downstream tasks. Nicheformer is pretrained on over 57 million dissociated and 53 million spatially resolved cells across 73 tissues from both human and mouse. Subsequently, the model is fine-tuned on spatial tasks for spatial omics data to decode spatially resolved cellular information.

We demonstrate the usefulness of Nicheformer in both linear-probing as well as fine-tuning scenarios on a novel set of spatially-relevant downstream tasks such as spatial density prediction or niche and region label prediction. In particular, we show that Nicheformer enables the prediction of the spatial context of dissociated cells, allowing the transfer of rich spatial information to scRNA-seq datasets. We define a series of novel spatial prediction problems and observe consistent top performance of Nicheformer, demonstrating the advantage of the improved model capacity of the underlying transformer. Additionally, we benchmarked Nicheformer in these tasks against scGPT^1^, Geneformer^2^, scVI^3^ and PCA and show that the Nicheformer architecture excels in these tasks. Altogether, our large-scale resource of more than 110 million cells in a partial spatial context, together with the set of novel spatial learning tasks and the Nicheformer model itself, will pave the way for the next generation of machine-learning models for spatial single-cell analysis.

## Introduction

Single-cell genomics technologies have advanced our understanding of cellular heterogeneity of entire tissues, organs, and organisms. Large-scale data generation efforts have led researchers to chart atlases of cellular heterogeneity of specific tissues and organs, such as the lung^4^, heart^5^ as well as broader cross-tissue atlases^6,7^. However, single-cell RNA sequencing technologies have the intrinsic limitation of requiring cells to be dissociated from their native tissue context. This results in a loss of information about the cellular microenvironment, which leads to an incomplete picture of the molecular factors of variation^8,9^. Recent advances in image-based spatial transcriptomics technologies have enabled the measurements of single-cell RNA-seq *in situ*, facilitating the profiling of hundreds of genes in hundreds of thousands of cells across various tissues and organs^9,10^. In situ spatial omics has been successfully applied to many biological systems, tissues and organs, revealing spatial components of cellular variations such as cell-cell communication^11^ and spatial gradients as well as emergent properties of tissue niches^12^, for example, in the mouse and human brain^13,14^, lung^15^, and liver^16^. Here we hypothesize that spatial omics data is becoming rich enough to learn a spatially aware, ‘foundational’ representation of cellular variation at scale.

A foundation model is a deep learning model trained on broad data that can be adapted to a wide range of downstream tasks^17^. In recent years, foundation models have emerged as a game-changing modeling paradigm, revolutionizing fields such as natural language processing^18,19^ and computer vision^20^. Foundation models increasingly account for multimodal data, by leveraging not only one data modality, e.g. text, but also images, video, and audio^21^. By leveraging massive datasets, powerful machine learning architectures, and large compute resources, foundation models are able to learn general representations of language, vision, or domain-specific data such as DNA^2,22,23^ and protein sequences^24–26^, outperforming more classical modeling approaches in a series of downstream tasks. Most commonly based on transformer architectures^27^, these models are pretrained on vast, unlabeled data, thus ditching the classical, costly tasks of sample annotation. Through self-supervision^28^, they learn powerful representations of complex data by identifying patterns and relationships within the data itself without the need for human-labeled examples. These learned representations then serve as a strong base for downstream tasks, while fine-tuning on labeled data further enhances performance on specific applications.

The field of single-cell biology has taken up deep-learning-based representation learning for some time, leveraging autoencoders^29–31^ for analysis tasks such as data integration^32^, atlas mapping^33^ and perturbation prediction^34–37^. Recently, the field has witnessed a surge in the development of foundation models designed explicitly for single-cell genomics data^1,2,23,38–45^. These models differ in tokenization and learning strategies, yet most of them leverage the transformer architecture with self-attention^27^. They rely on large data collections, usually in the order of tens of millions of cells, for pretraining^1,2,39^. The representations of genes and cells learned by these models are derived from implicitly modeling the complex interplay between gene expression patterns within a single cell via the flexible transformer architecture. Single-cell foundation models are evaluated on a variety of downstream tasks, such as cell type classification^2,39^, gene regulatory network inference^1,2^, or prediction of cellular responses to perturbations^1,41,43^. The diversity and complexity of these tasks are useful to thoroughly probe the model’s performance and to evaluate the robustness of the learned representation and the model’s ability to generalize to complex predictive tasks. Current results are promising but not entirely replicated in independent benchmarks^46–51^. Notably, to date, none of these models account for spatial relationships of cells during training, with the exception of CellPLM^41^, which, however, is trained on a limited dataset of 9 million dissociated and 2 million spatial transcriptomics cells^41^ and not fine-tuned on spatial tasks beyond gene imputation.

We propose Nicheformer, a foundation model pretrained on large-scale single-cell and spatial transcriptomics data to enable predictions for spatially dependent tasks with limited data availability. To learn spatial cellular representation at scale, we compiled the, to date, largest collection of single-cell and spatial transcriptomics datasets, spanning over 110 million cells from both human and mouse from 73 different organs and tissues. The Nicheformer data collection is the first large-scale pretraining corpus that contains not only dissociated single-cell data but also comprises over 53.83 million cells that were measured using image-based spatial technologies. By incorporating contextual information through modality, organism, and assay tokens, Nicheformer is able to learn a joint representation of single-cell and spatial genomics. We designed a set of novel downstream tasks showing that both fine-tuned Nicheformer and a linear probing model trained on the Nicheformer embedding systematically outperform existing foundation models, specifically Geneformer^2^ and scGPT^1^, pretrained on dissociated data alone and embedding models like scVI^29,30^ and PCA for these tasks. We demonstrate that Nicheformer accurately transfers the spatial context identified in spatial transcriptomics onto dissociated single-cell data, allowing users to enrich classical single-cell RNA-seq data with spatial context. This work paves the way for a new generation of foundation models for learning robust representations of cellular variation in tissues.

## Results

A transformer-based foundation model for combined spatial and disassociated single-cell data

### Overview

Nicheformer is a transformer-based model pretrained on a large curated transcriptomics corpus of dissociated and spatially resolved single-cell assays containing more than 110 million cells, which we refer to as SpatialCorpus-110M (Fig. 1A). Foundation model approaches build upon the idea of representing data in a way that allows a large language model, realized by the transformer architecture, to understand fine-grained dependencies in the data; the main differences lie in the way the data is encoded and how the model is trained and fine-tuned. Nicheformer’s novelty lies in generalizing previously employed tokenization strategies^2^ through the encoding of sample covariates across technologies modalities. This enables a general learning framework across modalities, opening up new possibilities for downstream tasks. We additionally enable learning multi-species embeddings with Nicheformer by defining orthologous genes across humans and mice (Methods), which was shown to work beneficial for cross-species biological investigations and enhanced the discovery of universal gene regulatory mechanisms^52^. We evaluated Nicheformer on a novel set of relevant downstream tasks that aim to demonstrate the model’s capability to transfer spatially inferred cellular variation to single-cell dissociated data (Fig. 1B).

**Figure 1.**
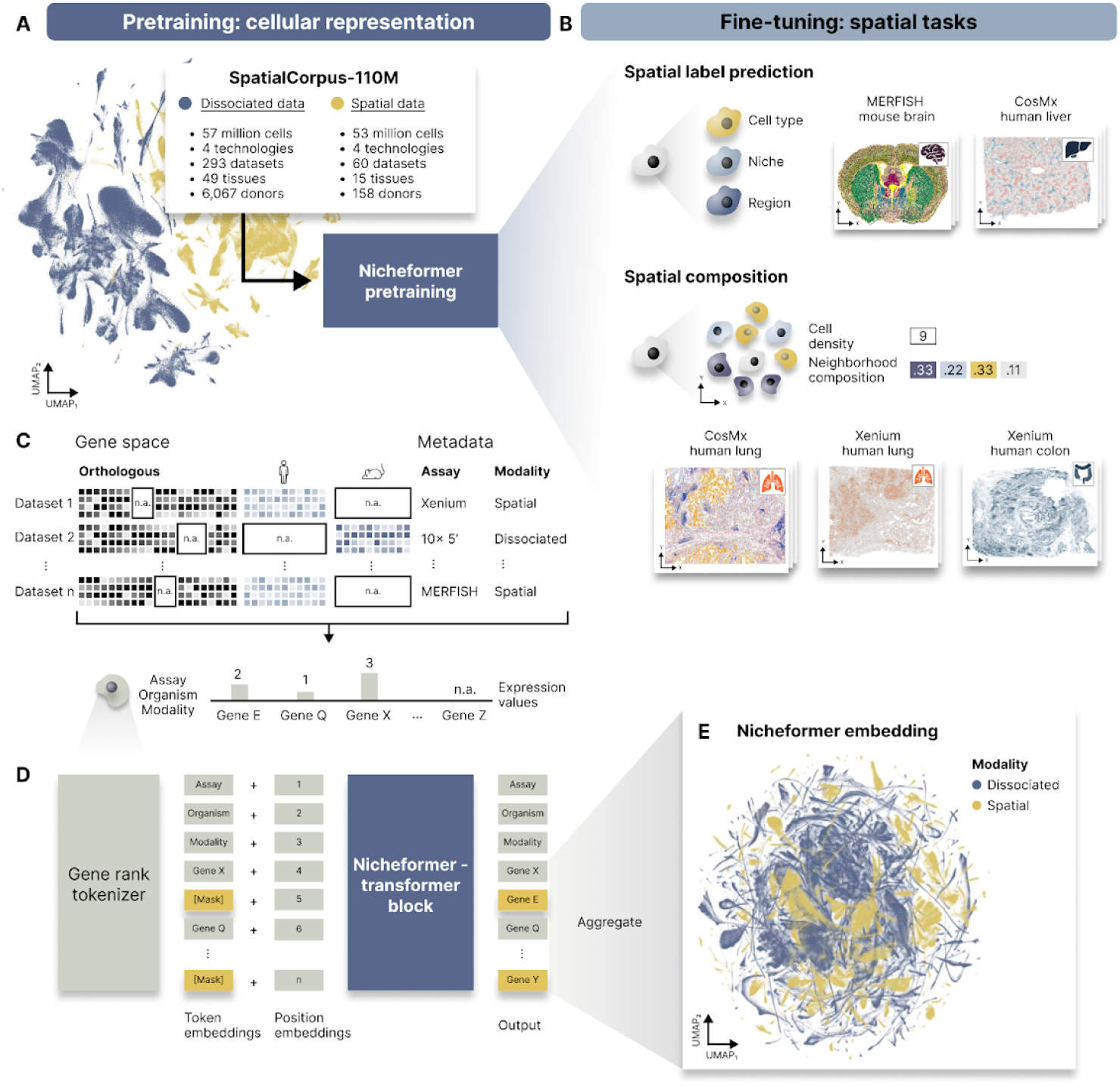
Nicheformer, a foundation model for spatial transcriptomics. A) Nicheformer is pretrained on the SpatialCorpus-110M, a large data collection of over 110 million cells measured with dissociated and image-based spatial transcriptomics technologies. The SpatialCorpus-110M collection comprises single-cell data from Homo Sapiens and Mus Musculus across 17 distinct organs, 18 cell lines, and additional single-cell data from other anatomical systems and junctions. Shown is an exemplary UMAP visualization of a random 1% subset of the entire pretraining dataset (n=1,108,759 cells) of the non-integrated log1p-transformed normalized SpatialCorpus-110M colored by modality. B) Nicheformer includes a novel set of downstream tasks, ranging from spatial cell type, niche and region label prediction to neighborhood cell density and neighborhood composition prediction. We test our approach on large-scale, high-quality spatial transcriptomics data from the brain (mouse - MERFISH), liver (CosMx - human), lung (CosMx - human, Xenium - human), and colon (Xenium - human). Visualized are example slices of the respective datasets colored by niche labels (brain, liver, and lung) and cell density (lung and colon). C) The SpatialCorpus-110M is harmonized and mapped to orthologous gene names, as well as human and mouse-specific genes, to create the input for Nicheformer pretraining. We harmonized metadata information across all datasets, capturing species, modality, and assay. D) Each cell’s gene expression profile and metadata are fed into a gene rank tokenizer to obtain a tokenized representation for each cell. The tokenized cells serve as input for the Nicheformer transformer block to predict masked tokens. Finally, the Nicheformer embedding is generated by aggregating the gene tokens (Methods). E) The pretrained Nicheformer embedding is visualized as UMAP colored by modality. The UMAP shows a random 5% subsample of the entire Nicheformer embedding (n=4,903,086).

The Nicheformer pretraining corpus comprises transcriptomics data from both humans and mice (Fig. 1A). Only expression data was used during pretraining to train the model to integrate data from dissociated and targeted spatial technologies. Both technologies exhibit substantial technical batch effects (Fig. 1A). A limiting factor for image-based spatial transcriptomics data is the targeted feature space which only measures a few hundred to a few thousands of genes, depending on technology and panel^53,54^. By pretraining across both modalities on the entire collection of datasets, instead of training organ-specific models, we aim to account for cross-tissue, cross-technology and cross-disease variations. To evaluate the representation learned in the Nicheformer pertaining, we focused for the downstream tasks on large-scale spatial transcriptomics datasets measured in four different solid organs with three image-based spatial transcriptomics technologies (Fig. 1B). We fine-tuned the pretrained Nicheformer model or performed linear probing on the pretrained Nicheformer embedding for each task and dataset separately. Linear probing consists of extracting the Nicheformer embeddings from the frozen transformer backbone and passing them through a single linear layer, designed either for regression or classification tasks, e.g. label prediction (Methods). The embedding is obtained by calculating a forward pass for a specific dataset through the pretraining model to generate a lower-dimensional representation, the so-called Nicheformer embedding. The organ-specific spatial context learned by the fine-tuned or linear probing Nicheformer model can then be used to evaluate the model’s ability to generalize information learned from spatial transcriptomics data, without directly accounting for the available spatial context, and transfer it to dissociated data.

### Cell representation

We define a cell as a sequence of gene expression tokens whose order represents the level of expression of each gene. Since the SpatialCorpus-110M is composed of two different species, we represent their expression profiles by concatenating matched orthologous genes using just the protein-coding ones as well as genes uniquely present in either of the two species, resulting in an overall vocabulary size of 20,310 gene tokens (Fig. 1C, Methods). To sequentialize a single-cell expression vector, we need to rank individual gene measures in relation to the mean gene expression in the SpatialCorpus-110M. This way, each cell is converted into a sequence where each token is a gene, further referred to as a gene token, and those are ordered according to its relative expression (Fig. 1D, Methods). Encoding cells based on their gene ranks has been shown to generate embeddings that are robust to technical batch effects but preserve the gene-gene relationships in a cell^2^. We first concatenate all technology-specific datasets from the SpatialCorpus-110M and pad missing genes not measured for individual cells. Previous works^54,55^ have demonstrably shown technology-dependent biases between spatial and dissociated transcriptomics data. Since spatial transcriptomics measurements and the obtained signal heavily depend on the chosen preprocessing pipeline used to generate cell-by-gene count matrices^56^, average expression profiles obtained with image-based spatial transcriptomics technologies can show much higher gene counts compared to dissociated technologies^54^. We argue that instead of calculating a non-zero mean vector across the entire SpatialCorpus-110M, it is preferred to compute a technology-specific non-zero mean vector, which accounts better for assay-specific gene expression effects. We therefore grouped all dissociated assays into one technology, whereas for the spatial transcriptomics datasets, we defined the overall technology categories as MERFISH, Xenium, CosMx and ISS. The technology-specific non-zero mean vector is then given by the mean expression value of each gene, without considering the zero counts. We additionally account for different modalities, technologies and species by adding contextual tokens that model the nuances of each of those elements. By incorporating the contextual tokens, the model is able to learn the nuances that differentiate organisms, modalities, and technologies in the measured cells.

We evaluated whether Nicheformer is robust to the commonly used gene ranking strategy^2^, in particular when a subset of the genes is not used as input to the model. This evaluation resembles the real-world challenge of probe-set selection in spatial transcriptomics, where only a small set of genes is measured. We randomly shifted, removed or transposed the ranks of individual or groups of genes (Methods). Next we embedded the cells with perturbed ranks with Nicheformer and used integration metrics^32^ to assess whether the non-perturbed-rank and perturbed-rank cells are embedded closely by the model. We observed that cell embeddings are stable in both technologies up to a perturbation of 20 % in their gene ranks in terms of silhouette score (Suppl. Fig. 1). This analysis shows that Nicheformer’s embeddings are sufficiently robust to input perturbations such as incomplete gene panels.

### Model design and training

The Nicheformer architecture uses a context length of 1,500 gene tokens serving as input for its transformer. The transformer block leverages 12 transformer encoder units^19,27^ with 16 attention heads per layer and a feed-forward network size of 1,024 to generate a 512-dimensional embedding of the pretraining dataset, resulting in altogether 49.3M parameters, which resulted in the best performance compared to models with fewer parameters (Suppl. Fig. 2) and other model hyperparameters (Suppl. Table 1). Nicheformer uses self-attention to weigh the importance of each gene in a cell in relation to every other gene, enabling it to capture gene-gene relationships effectively throughout pretraining. We confirmed technology-dependent biases between spatial and dissociated transcriptomics data by evaluating initial pretraining experiments that were trained on just a single modality (Suppl. Fig. 3). We found that only training on dissociated single-cell data resulted in a lower performance in terms of F1 score for the task of annotating cells in a spatial transcriptomics dataset in the brain. This indicates that dissociated data alone can not resolve the sources of variation observed in spatial transcriptomics data.

### Model evaluation and downstream tasks

Current transformer-based models designed for single-cell genomics data are used for either gene-level tasks, such as inference of gene regulatory networks and prediction of perturbation effects, or cell-level tasks, such as cell type annotation and batch integration^1,2,39^. By incorporating single-cell data at both the dissociated and the spatial scale into a single model, we are able to define a novel set of biologically meaningful spatial tasks, where previous models primarily only focused on disassociated ones (Suppl. Table 2). These spatial tasks formulate biological questions related to the cellular composition of tissues and are sufficiently challenging to resolve: they address the prediction of human-annotated niche and region labels and spatial composition prediction (Fig. 1B, Methods). For the spatial label prediction tasks, we also evaluate the model’s uncertainty regarding the predicted labels (Methods). For spatial composition tasks, we define a distance-based spatially homogeneous niche around each cell and ask the model to predict local density or cell type composition. The tasks are formulated as prediction problems operating on Nicheformers pretrained embedding (Fig. 1E). Importantly, the obtained embedding is not comparable to commonly known integrated spaces as retrieved in single-cell atlases, but rather captures a cross-technology, cross-tissue and cross-species cell representation that reflects meaningful information usable for downstream tasks.

### Model transfer-learning

We investigate the performance of Nicheformer in a linear probing as well as a fine-tuning setting. Both fine-tuning and linear probing involve adding a new linear layer on top of the pretrained Nicheformer model adapted to the prediction task formulated above. Fine-tuning also updates the parameters within the Nicheformer model itself, while linear probing keeps those parameters frozen. Through the low complexity of the linear probing mode, we ensure that the biological insights result from the meaningful biological information captured by the learned Nicheformer embedding (Fig. 1E).

#### SpatialCorpus-110M, a large-scale, cross-organ and cross-species pretraining dataset for single-cell and spatially resolved transcriptomics

To train the Nicheformer model, we collected and harmonized a new, large-scale pretraining corpus of single-cell and spatially-resolved transcriptomics data from human and mouse tissues. To our knowledge, this resource consists of the, to date, largest collection of harmonized and combined single-cell and spatially resolved transcriptomics datasets.

SpatialCorpus-110M consists of a total of 57.06 million cells from single-cell dissociated technologies. It builds upon the CellXGene CENSUS database (33.47 million cells), which we extended by an additional 180 datasets across 73 different tissues, containing 17 solid organs, 18 cell lines and various additional tissue junctions in human and mice, with harmonized ontologies and metadata (Fig. 2A). These additional dissociated datasets have been collected through Gene Expression Omnibus^57,58^, sfaira^59^ and the Human Cell Atlas data explorer^60^ (Suppl. Table 3, Methods).

**Figure 2.**
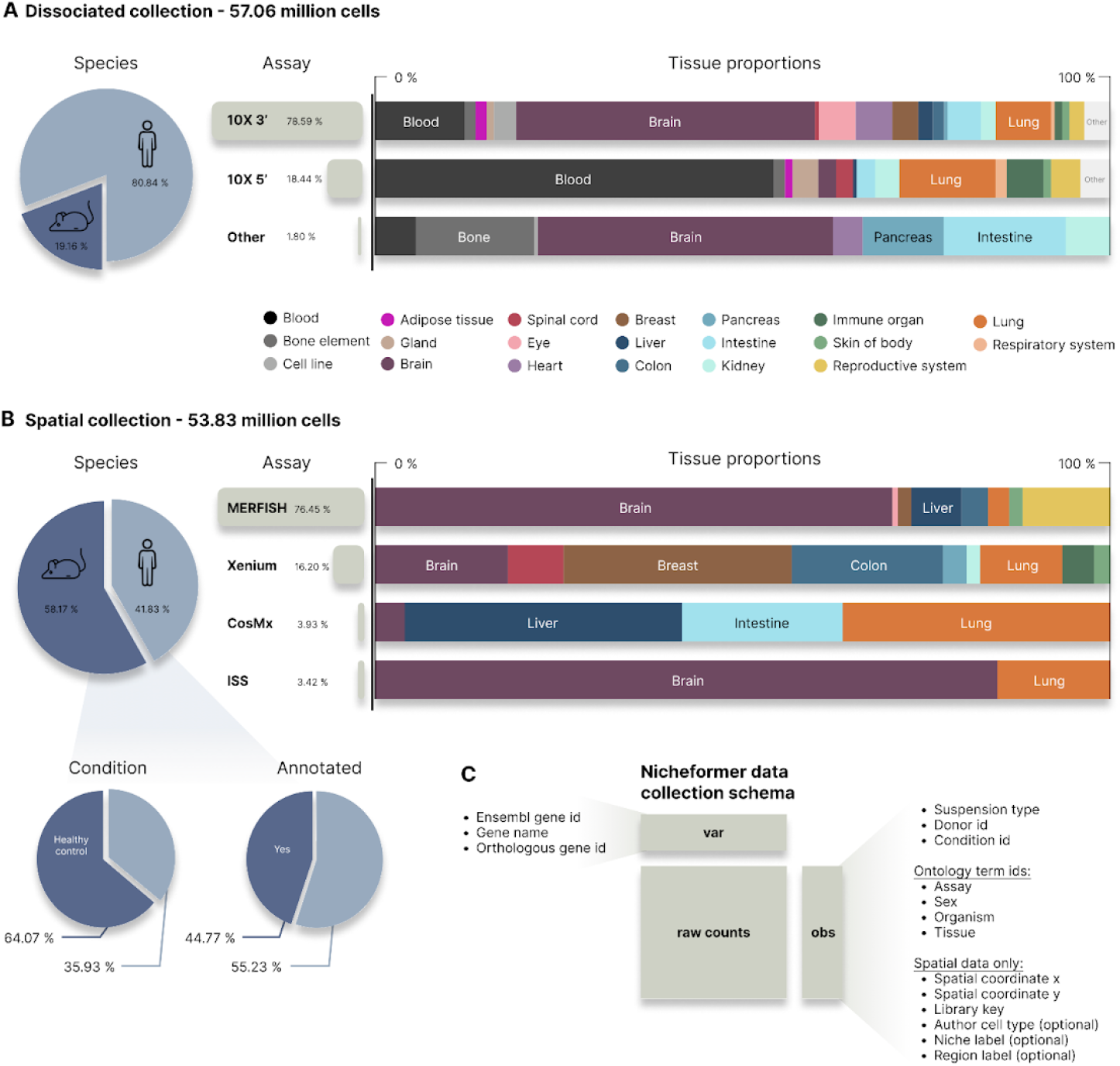
Overview of the SpatialCorpus-110M collected for training Nicheformer. A) The dissociated single-cell genomics data collection contains 57.06 million human and mouse cells. The collection includes cells from 17 different organs, 18 different cell lines, blood, bone elements, tissue junctions, and other anatomical entities, grouped by primary solid organ to simplify visualization and analysis. B) The spatial transcriptomics data collection contains 53.83 million targeted spatially-resolved cells obtained from humans and mice. The collection comprises four different profiling technologies across 15 different solid organs. C) The SpatialCollection-110M was collected with harmonized metadata defined in the Nicheformer data collection schema (Methods). Metadata was harmonized both on gene and on cell level depending on modality.

Altogether, the dissociated collection of SpatialCorpus-110M comprises cells from over six thousand different donors and technical or biological replicates. For spatial transcriptomics, we curated image-based spatial datasets, specifically MERFISH^61^ (Vizgen MERSCOPE), 10x Genomics Xenium, Nanostring CosMx^15^, and In Situ Sequencing (ISS)^62^ data (Fig. 2B, Suppl. Table 4). Data was collected from individual publications as well as via the Vizgen data release^63^ (18.8%) and the 10x Genomics data resource^64^ (13.7%). Altogether, the spatial data consists of 53.8 million cells across 15 tissues in human and mouse, including 158 individuals or animals and over 10,600 individual tissue sections. The biggest proportion of cells in the spatial collection originated from the brain, with 60.46% (n=32,146,779 cells) of all cells collected, followed by the lung with 9.95% (n=3,199,548 cells). A large proportion of the publicly available spatial omics datasets we collected is not annotated (55.23%). Cells are mostly from healthy controls (64.07%), but we also included cells from various cancer samples (31.98%) to enable Nicheformer to learn meaningful representations of the tumor-immune microenvironment in space.

For all datasets in the Spatial-Corpus-110M, we curated metadata, such as assay, sex, organism, and tissue, based on the original publications by using official ontology term identifiers (Fig. 2C, Methods). To harmonize features across species, tissues and assays, we first converted all gene symbols to ENSEMBL gene IDs using pyEnsemble^65^. Then we used BioMart^66^ through the official Ensembl releases^67^ to match orthologous genes between species. This resulted in 16,981 orthologous genes matched between mouse and human, 151 genes specific only to mouse and 3,178 genes specific only to human. This harmonization generated a vocabulary size of 20,310 unique gene ID tokens.

#### Nicheformer learns sex-related difference in gene-gene dependencies in MERFISH mouse brain data

Understanding differences across conditions is a crucial and important analysis step for single-cell data. By learning on large data collections, foundation models learn sources of biological variation in terms of gene-gene relationships that can provide meaningful access points for differential single-cell analysis. We evaluated whether Nicheformer pretraining and obtained attention matrices capture meaningful biological variations in two MERFISH mouse brain datasets that were obtained from a male and a female mouse respectively^13^ and are part of the SpatialCorpus-110M (Methods). Both datasets are are annotated with the same set of annotations and therefore allowed us to performed a targeted analysis of those by leveraging the common coordinate framework (CCFv3) annotations provided by the Allen Brain Institute. Specifically, the anteroventral periventricular nucleus (AVPV) is known to exhibit sex-dependent morphology and gene expression^68^. During pretraining, Nicheformer encodes male and female cells without any covariate annotation. Hence, it is the internal attention mechanisms that implicitly learn to differentiate between male and female biological signatures. These mechanisms allow the model to capture subtle differences in gene expression and other biological patterns related to sex, even in the absence of explicit labeling. To understand and delineate such mechanisms, we analyzed the attention matrices of 2,000 cells from both male and female subjects from the AVPV section and uncovered distinct patterns across the attention heads (Fig. 3A-C).

**Figure 3.**
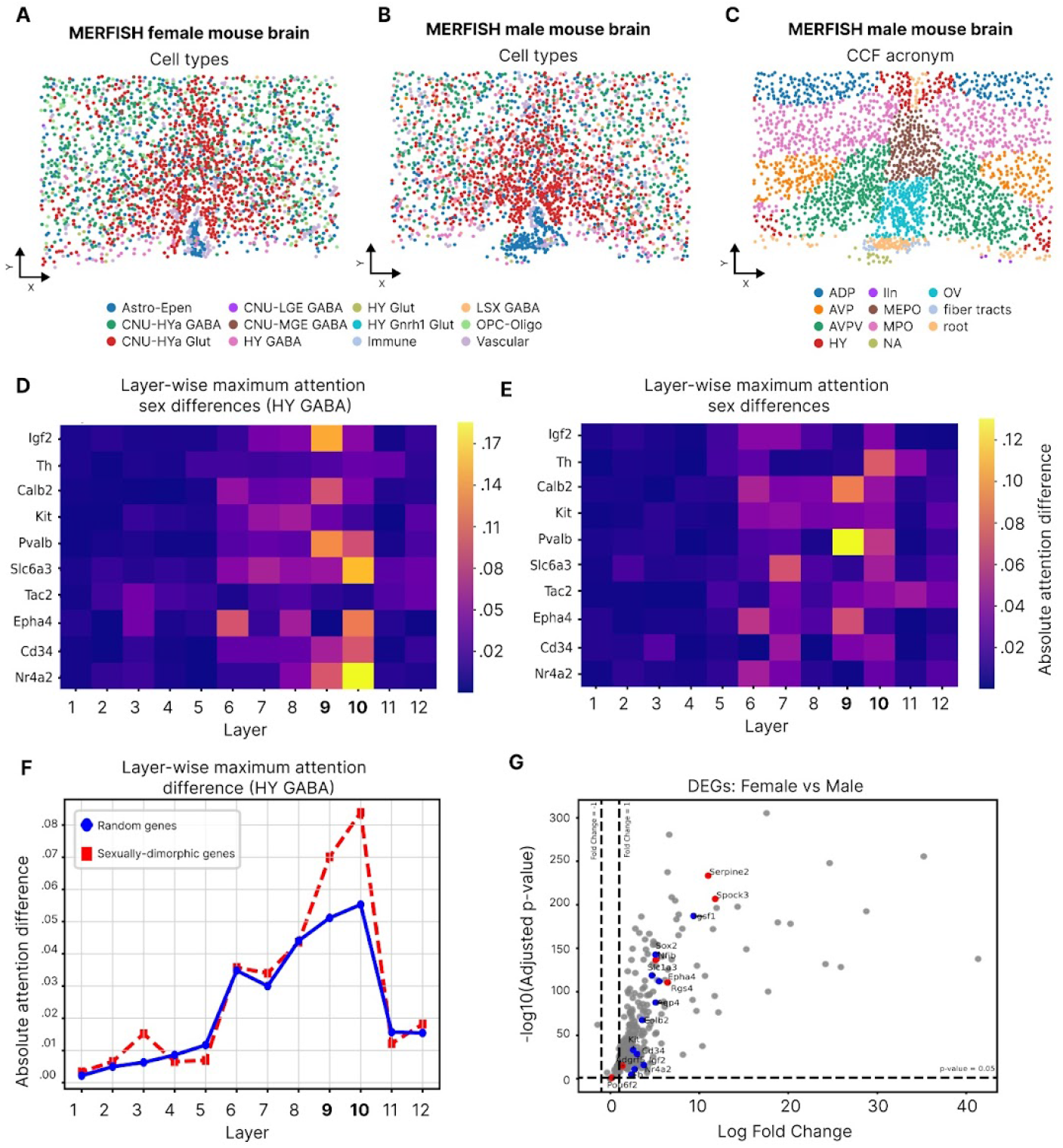
Nicheformer identifies gene-gene dependencies between male and female MERFISH mouse brain sections. A) Single-cells resolved in space on an example slice (n=2,292 cells) of the MERFISH female mouse brain dataset with cell type label superimposed. B-C) Single-cells resolved in space on an example slice (n=2,269 cells) of the MERFISH male mouse brain dataset with the cell type label (B) and CCF acronym label label (C) superimposed. D-E) Absolute difference of layer-wise attention scores between male and female MERFISH mouse brain sections show per transformer block of the sexually-dimorphic genes (SDGs) considering just the HY GABA cells (D) and the entire AVPV section (E). F) Maximum layer-wise attention difference across layers between male and female HY GABA cells. It can be seen that the attention payed to random and SDGs is equal across all layers except in layers 9 and 10, where there is an increment in the maximum attention payed to the SDGs in comparison to the attention payed to the random set of genes G) Volcano plot showing the differentially expressed genes highlighting the genes with highest attention difference between sexes (red), and highlighting the SDGs (blue). It can be seen that the genes found with highest attention differences are not among the most differentially expressed.

Some attention heads exhibited a strong focus on metadata tokens, while others distributed their attention more evenly across the entire context length. Additionally, we identified attention heads with highly characteristic patterns, that range from a strong self-attention focus (visualize as strong diagonal attention scores) to off-diagonal patterns that indicate that the attention is distributed across genes with similar normalized expression (and therefore closer in the input sequence) (Suppl. Fig. 4). These findings highlight the diverse range of attention behaviors that Nicheformer develops when processing complex biological data. Notably, metadata tokens were generally highly attended, with the last transformer blocks demonstrating a pronounced focus on these tokens, suggesting their significance in the overall learning process (Suppl. Fig. 5).

We additionally compared the models learnt attention for a predefined set of ten genes from the original 500-plex gene set of both MERFISH datasets in the same ∼2000-cell squares of both a female and male sections containing the AVPV. These genes were selected based on previously reported sexually-dimorphic expression^69–74^. Next, we extracted the all attention matrices from Nicheformer’s 12 transformer layers for each cell in the AVPV section of both male and female mice and compared the attention paid to the predefined set of genes. HY GABA neurons are a small population of cells in the AVPV which modulate the firing of the different glutamatergic neurons in the AVPV which stimulate the synthesis of gonadotropins^75^. We do the analysis for both all cells in the AVPV section and just HY GABA cells, to get a fine grain detail of a specific cell type in which we expect sex differences in the AVPV section. We identify key differences between the male and female cells (Fig. 3D-E). The first eight layers had the greatest average attention differences for both sexually-dimorphic genes (SDGs) and 100 random genes not directly linked to sex-specific differences in the brain (Suppl. Fig. 6A-B). In contrast, layers nine and ten show high maximal attention value differences for SDGs, when performing differential testing on the attention weights between those two groups, specially for the HY GABA cells (Fig. 3D-F). This suggests that there are noticeable differences in the attention paid to the SDGs between male and female subjects across those layers. Interestingly, the contrast between the average and the maximum attention difference indicates that the sex differences are captured by a subset of the attention heads, with at least one of the 16 attention heads showing a stronger focus. Importantly, this contrast between the average and the maximum difference in attention also holds for genes in the random set (Suppl. Fig. 6C-D). Furthermore, six out of the ten genes with the highest attention differences between sexes (*Adgrf5, Nfib, Pou6f2, Rgs4, Serpine2, Spock3*) have not previously been reported to have sexually dimorphic expression in the brain and some were not differentially expressed between the male and female brain section (Fig. 3G). These genes are either transcription factors involved in neural development, regulate g-protein coupled receptors (GPCRs), or extracellular matrix (Serpine2, Spock3), all functions where we expect sex differences in the AVPV, however, none of these genes have known sexually dimorphic expression or function. This is in part a limitation of the gene panel, which was designed to differentiate cell types rather than sex differences. Instead, sex differences that are encoded in the interactions of these non-sexually dimorphic genes with both the measured SDGs and other, unmeasured genes normally used to demarcate sex differences in the AVPV like kisspeptin/receptor (Kiss1/Kiss1r), gonadotropin releasing hormone (Gnrh), or estrogen receptor alpha (Esr1)^76^, are uncovered through the attention mechanisms of Nicheformer. Some, but not all the genes with high attention differences are differentially expressed between the male and female brains, although this is not a great measure as the female brain has much higher counts overall so most of the gene panel have higher expression in the female gene panel. This highlights one strength of Nicheformer: thanks to its ranked based tokenization and the attention mechanisms, it is not reliant on perturbation datasets necessarily having comparable measurement depth.

#### Nicheformer allows transferring spatially-resolved cell type, niche and region labels onto unseen data

Dissociated single-cell atlases excel at capturing molecular maps of the cell type diversity of tissues and organs^4^. Cells are typically well described by their cell type, commonly defined as a group of cells that share a characteristic and stable molecular state, reflected in their structure and function^77^. These cell types can be found in one specific tissue or across multiple tissues^7^. Typically, cell types are defined without directly incorporating the cellular microenvironment^78^. However, a cell’s spatial context provides additional value and information beyond the cell’s expression profile and can be helpful for understanding cellular microenvironments^79^. Spatially-resolved single-cell genomics allows us to augment the concept of cell types when considering also the molecular state of the neighbors of a cell. We refer to this as a cell niche. In contrast to standard approaches to identifying cell types, cellular niches are defined by spatially-dependent labels obtained by combining similarity in gene expression profiles together with spatial neighborhood structures, histological information or global structures in the tissue^12,80–82^. Cellular niches typically describe a local structure in a tissue, for example, a tumor niche or immune niche, which vary both in terms of gene expression, but also in terms of the cell type compositions. Similarly to niches, tissue regions can be defined as spatially consistent cellular states across a larger-than-a-niche tissue context. Importantly tissue regions are commonly composed of multiple different niches and therefore describe higher-level structures in a tissue.

Due to the technical limitations of spatial omics data, transferring labels from dissociated single-cell data, and vice versa, is often challenging as it requires well-suited reference datasets with a high fraction of overlapping genes between both^83^. As the feature space of targeted spatial transcriptomics data is still limited, methods for reference mapping are also trained only on a limited set of features and therefore often perform poorly. Additionally, methods typically are designed for the use-case, one spatial omics dataset and one reference atlas, but they are not designed to learn from reference atlases at the scale of hundreds of million cells. Nicheformer enables researchers to leverage a large scale reference atlas through the SpatialCorpus-110M for mapping annotations between both modalities and therefore addresses existing limitations present for this task.

To test whether Nicheformer, pretrained only on gene expression information, can efficiently transfer spatial context with minimal to no supervision we considered a large MERFISH mouse brain dataset^13^, where 17 different brain regions were defined by registering a 10×10 µm grid of cells assigned to their anatomical parcellation of the Allen Mouse Brain Common Coordinate Framework^84^ v3 (Suppl. Fig. 7A). The resulting CCFv3 registration was then used to obtain a region label for each cell measured in the MERFISH mouse brain dataset^13^. This dataset additionally consists of seven distinct tissue niches (Fig. 3A) defined based on their molecular and anatomical similarity, which are structurally organized across the entire brain (Suppl. Fig. 7B).

In this setting, we assessed Nicheformer’s ability to accurately impute niche and region labels on unseen, held-out tissue sections from the MERFISH mouse brain dataset, measuring one male mouse brain (Suppl. Fig. 7A-D). We evaluated two versions of Nicheformer, a direct use of the pretrained embeddings (Suppl. Fig. 8A) via linear probing as well as fine-tuning the model for the prediction task. To assess model performance, we ask whether the Nicheformer embedding better captures information required for the downstream niche and region prediction task compared to more classical embeddings calculated with a variational autoencoder (scVI) and PCA on the specific data set without pretraining - in disassociated tasks it has proven to difficult for foundation models to outperform data set specific embeddings. First, we trained scVI and PCA on the training set of the MERFISH mouse brain dataset and then generated an embedding for all cells present in the dataset (Methods). We then used the resulting embeddings as input for a linear classifier in a similar way as done for the Nicheformer representation (linear probing). Additionally, we also train PCA and scVI in a subset of SpatialCorpus-110M to assess whether a larger training set would improve their results and repeat the same linear probing procedure. Finally, we also evaluate the linear probing scenario for foundation models trained on dissociated data alone: GeneFormer and scGPT.

We found that both of our approaches outperform both traditional embedding methods, scVI and PCA, no matter which was the training set, as well as GeneFormer and scGPT, in terms of macro F1 score, which effectively balances precision and recall across all classes (Fig. 4B). Notably, while the linear probing approach consistently outperforms the baselines, fine-tuning Nicheformer further enhances performance, demonstrating the capability of the model pretrained only on gene expression information to adapt to spatial tasks. We performed a similar analysis on a randomly held-out test set of the CosMx human liver dataset defining tissue niches as different zonations between the central and portal veins (Suppl. Fig. 9A-C). Again, the fine-tuned Nicheformer model showed the best performance in terms of macro F1 score. However, the Nicheformer linear probing model did not outperform linear probing models trained on scVI and PCA models - trained on the training set of the liver dataset (Suppl. Fig. 9F). We hypothesized that the worse linear probing performance is related to the insufficient model capacity due to limitations in the number of trainable parameters and training time. This prevents the model from accurately learning a nuanced representation of liver cells, which have relatively low overall abundance in the SpatialCorpus-110M (Fig. 2A, B). We, therefore, asked whether additional pretraining of Nicheformer solely on the liver training set improved performance. Indeed, we observed an improvement in linear probing performance for the prediction of the liver niche labels on the longer pretrained Nicheformer embedding (Suppl. Fig. 9F). We conclude that longer pretraining can be beneficial for datasets with lower abundance in the training corpus.

**Figure 4.**
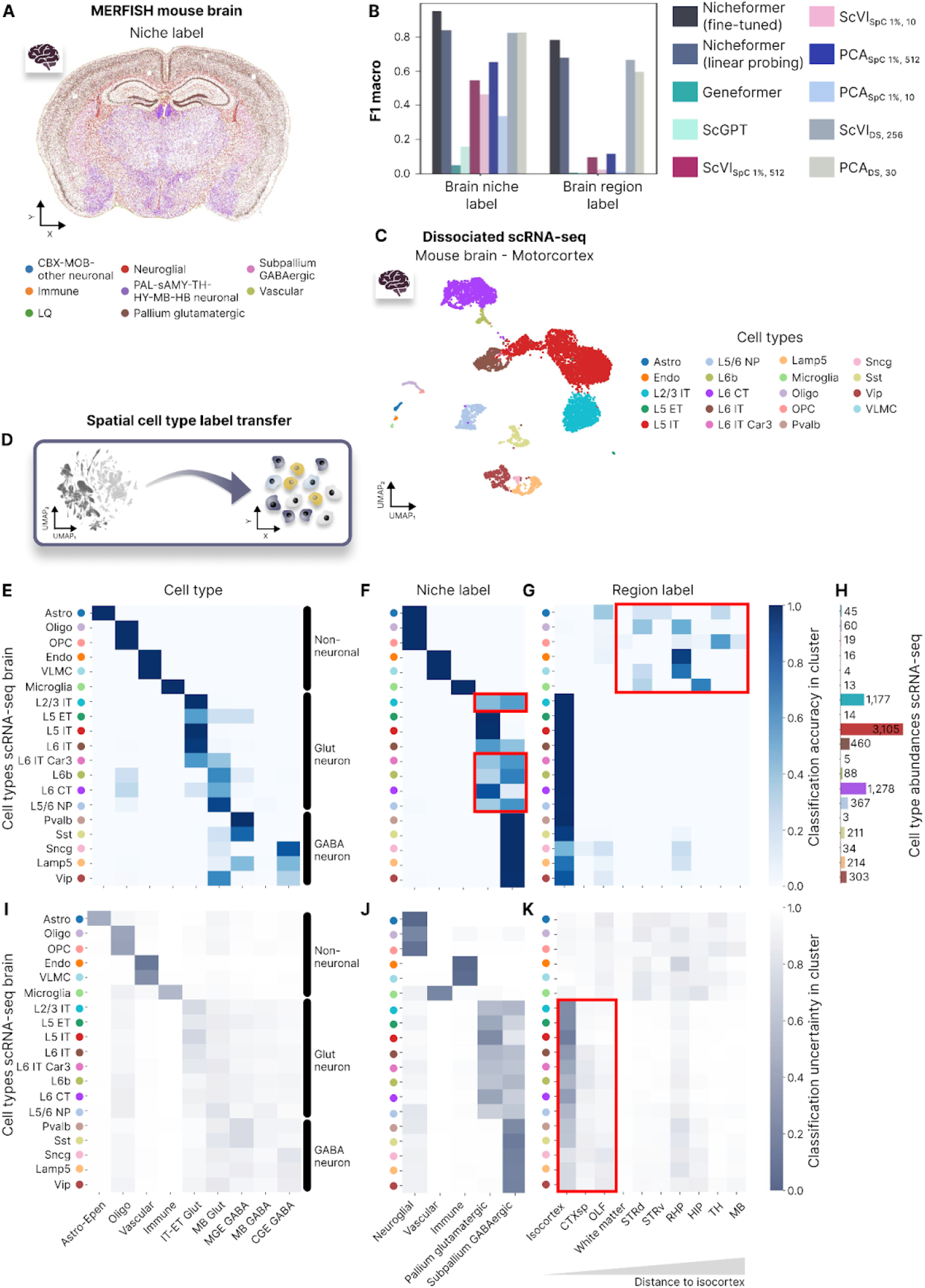
Nicheformer accurately transfers cell type, niche, and region label to unseen spatial and dissociated data in the brain. A) Single-cells resolved in space on an example slice (n=114,396 cells) of the MERFISH mouse brain dataset with niche label superimposed. B) Test-set F1-macro of niche and brain region label prediction of the fine-tuned Nicheformer model, the linear probing model, and a linear probing baseline computed based on embeddings generated with Geneformer, scGPT, scVI and PCA, respectively. For scVI and PCA, both embeddings generated from a random 1% subset of the SpatialCorpus as well as embeddings generated from the training set of the original dataset are evaluated. C) UMAP of dissociated scRNAseq dataset with original author cell type label superimposed. D) Nicheformer can transfer spatial niche and region labels onto dissociated single-cell data. E) Nicheformer accurately classifies cells from the dissociated motor cortex to relevant cell types (n=9 out of 33 distinct ones in the classifier) trained on the whole mouse brain MERFISH dataset. F-G) Nicheformer correctly projects dissociated single-cells to niche (F) and region (G) labels to provide spatially dependent labels. F) Nicheformer misclassified parts of L2/3 IT neurons as residing in the subpallium GABAergic niche (highlighted in the red box). Additionally, the deep cortical excitatory neurons L6b, L6 CT, L6 IT, and L6 IT Car3 (highlighted in the red box) should be classified as pallium glutamatergic niche instead of subpallium GABAergic by Nicheformer. G) Most of the non-neuronal cells (84.7 % of all non-neuronal cells, n=133) were misclassified as not belonging to the isocortex or the adjacent brain regions (highlighted in the red box). H) Cell type abundances in the scRNA-seq dataset measuring the primary motor cortex in the mouse. I-K) Classification uncertainty of label transfer of the dissociated scRNA-seq dataset to the MERFISH mouse brain data for cell type label (I), niche label (J), and region label (K) with a value of 0 representing a high uncertainty and 1 being a lower uncertainty, i.e., high certainty. K) Observed high uncertainty for parts of the Glut and GABA neurons for the region prediction of the isocortex, CTXsp and OLF, which are neighboring brain regions.

Next, we assessed whether Nicheformer accurately transfers cell type labels defined in the MERFISH mouse brain dataset onto dissociated cells present in the SpatialCorpus-110M and measured in the primary motor cortex cells with scRNA-seq (Fig. 4C, D)^14^. We find that Nicheformer correctly selects the nine motor-cortex related cell types out of the overall 33 cell types present in the MERFISH mouse brain dataset (Fig. 4E, Suppl. Fig. 10A). When calculating classification uncertainty based on the overall predicted distribution generated by the model (Methods), the predicted cell type labels show overall a high agreement and low classification uncertainty (Fig. 4E, I) with the original cell type annotations. Nicheformer correctly assigns all non-neuronal cells to their mapped spatial cell type (Fig. 4E, Methods). Glutamatergic (Glut) neurons, which are the most abundant cell types in the dissociated dataset (Fig. 4H), are also correctly mapped to the correct neuronal identity (Glut); however, the regionalization for a few Glut neuronal subtypes from deeper cortical layers (L6b, L6 CT, L5/6 NP) is mispredicted as midbrain glutamatergic (MB Glut), while one would expect an assignment to NP-CT-L6b Glut (Suppl. Fig. 10A). The misclassification of the region of the glutamatergic neurons can be linked to the overall transcriptional diversity in the fine-tuning MERFISH mouse brain dataset. MB Glut cells are sub-clustered into 657 subtypes, whereas NP-CT-L6b Glut cells only define 83 distinct subtypes even though they are overall more abundant in the MERFISH mouse brain dataset (MB Glut - n=88,169 cells, NP-CT-L6b Glut - n=174,616 cells)^13^.

Next, we wanted to assess whether Nicheformer is able to annotate brain niche labels in the same scRNA-seq primary motor cortex dataset. We find that Nicheformer correctly predicts only the expected niche labels for all non-neuronal cell types with low uncertainty (Fig. 4F, J, Suppl. Fig. 10B). Additionally, all GABAergic (GABA) and Glutamatergic (Glut) neurons are correctly predicted to be part of either the pallium glutamatergic or subpallium GABAergic (Fig. 4F), yet with higher uncertainty for excitatory neuronal subtypes (Fig. 4J). However, we also observe a partial misclassification for parts of the Glut neurons, which is likely due to the spatial overlap of the pallium glutamatergic and subpallium GABAergic niches (Suppl. Fig. 10D, E) and the high diversity of cell types within the pallium glutamatergic and GABAergic niches.

We then assessed whether Nicheformer correctly annotates brain regions to the primary motor cortex dissociated cells. The expected region for all cells is the isocortex. Nicheformer correctly predicts the isocortex for all neurons, with very low uncertainty for the excitatory neurons and slightly higher uncertainty for the inhibitory neurons (Fig. 4G, K, Suppl. Fig. 10C). We also observe that some cells are classified in brain regions neighboring the isocortex, namely the cortical subplate (CTXsp), olfactory area (OLF), and white matter (Fig. 4G, K). This classification might be plausible as during tissue dissection other brain regions might be additionally captured. For region classification, we observed a lower performance compared to cell type and niche label for the non-neuronal cells. This could be related to the lower transcriptional diversity and their lower regional specificity compared to neuronal cells^13^.

Altogether, this demonstrates Nicheformer’s ability to learn powerful cell representations by capturing nuanced spatial information. Linear probing on the pretrained model already surpasses existing baselines, highlighting the effectiveness of the representation. Fine-tuning further refines this representation, emphasizing the importance of task-specific adaptation for capturing subtle cellular variations. Notably, Nicheformer enables the direct transfer of spatially-aware annotations from spatial to dissociated single-cell data by using a simple linear layer. This capability unlocks new possibilities for analyzing single-cell data across different modalities.

#### Nicheformer predicts neighborhood compositions in spatial and dissociated single-cell data

Tissue microenvironments consist of cellular neighborhoods with a diverse composition of cell types. Differences in neighborhood composition have been shown to have a significant effect on gene expression and can be associated with cell-cell communication events^11^. Furthermore, the cellular composition of neighborhoods in the tumor microenvironment may hold prognostic value^85,86^, since immune cell infiltration in the spatial context is a predictor for cancer survival^87,88^. Here we show that we can leverage Nicheformer’s multimodal cell representation to accurately relate changes in gene expression to differences in neighborhood compositions in spatial data and transfer them to dissociated transcriptomes.

We define a cell’s ‘computational’ neighborhood as the set of cells within a fixed radius (Fig. 5A, Methods). The total number of cells composing the neighborhood defines the neighborhood density, and the proportion of cell types in the neighborhood defines the neighborhood composition. This notion is consistent with previous approaches defining a cellular neighborhood^89^ and allows for an interpretable evaluation of model results. Generally, the definition of a cell neighborhood can be extended in the future to account for non-isotropic cell neighborhoods that might vary in their cell type composition and are drivers of similar biological functions with varying sizes across a dataset.

**Figure 5.**
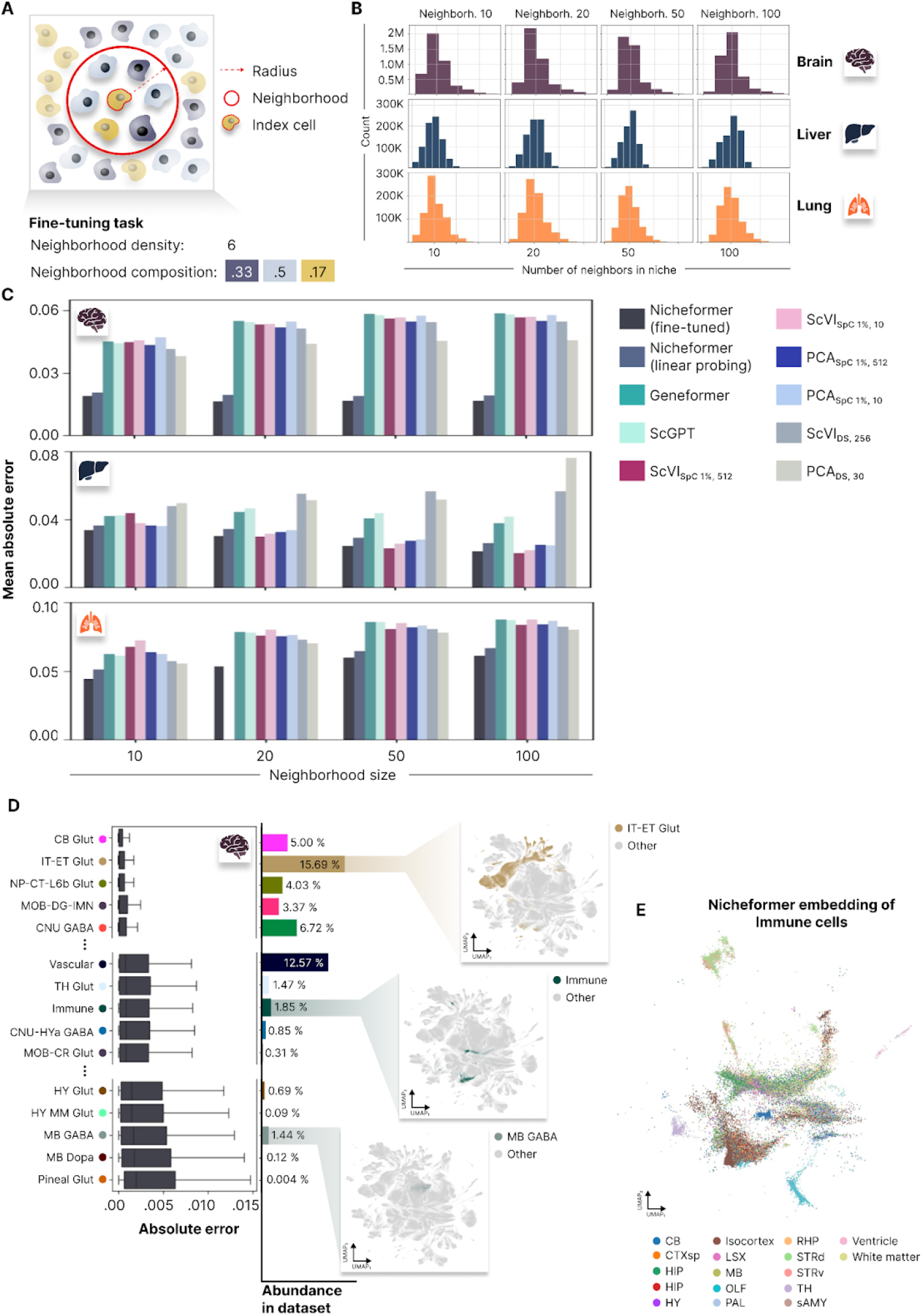
Nicheformer accurately predicts neighborhood compositions at multiple niche resolutions for the brain, liver, and lung. A) We define the neighborhood of a cell as its local neighborhood given a radius and an index cell. The neighborhood cell density is then defined by the number of cells in the neighborhood, and the neighborhood compositions are the proportions of neighboring cell types. B) Neighborhoods are computed at multiple resolutions resulting in different neighborhood size distributions. Each barplot shows the distribution of the number of neighbors across the brain, liver, and lung datasets. We extract neighborhoods with the mean number of neighbors 10, 20, 50, and 100 for each dataset. C) The fine-tuned and linear probing Nicheformer models outperform for brain and lung linear-probing models trained on Geneformer, scGPT, scVI and PCA embeddings in terms of mean absolute error across all neighborhood sizes. Still, it struggles to outperform all benchmarks in liver, where scVI models are very competitive. This is an issue related to the previous liver performance reported in the previous section (Suppl. Fig. 6F). D) Left: Fine-tuned Nicheformer performance on the MERFISH mouse brain data grouped by index cell type. Shown are the absolute error values between predicted and observed neighborhood composition vectors for held-out test cells. For each box in (D), the centerline defines the median, the height of the box is given by the interquartile range (IQR), the whiskers are given by 1.5 × IQR, and outliers are given as points beyond the minimum or maximum whisker. Center: Index cell type abundances in the entire MERFISH mouse brain dataset. Right: UMAPs of MERFISH mouse brain Nicheformer embedding with the selected index cell type as color superimposed. E) UMAP of the Nicheformer embedding of all immune cells in the MERFISH mouse brain dataset with region label as color superimposed.

To evaluate Nicheformer’s ability to predict neighborhood composition, we focused on three datasets measuring three organs with two different technologies, namely MERFISH mouse brain, CosMx human liver, and CosMx human lung. We computed neighborhood compositions at varying resolutions for each of the three datasets separately. The radii were selected to contain, on average 10, 20, 50, or 100 neighbors (Fig. 5B, Methods). We evaluated Nicheformer both in linear probing and fine-tuned settings for each dataset and each neighborhood size individually and compared its performance to linear probing on embeddings computed with scVI, PCA, Geneformer and scGPT. We found that fine-tuned Nicheformer systematically outperforms the linear probing models trained on Nicheformer embedding, Geneformer, scGPT, scVI and PCA for this task on all three organs in terms of mean absolute error. Notably, the linear probing models trained on Nicheformer embeddings also outperform all other methods, except for the fine-tuned Nicheformer. (Fig. 5C). However, for bigger radius sizes in the liver dataset, the scVI models trained in a subset of SpatialCorpus-110M perform on par with fine-tuned Nicheformer. We believe this to be related with the previous classification results in the same dataset (Suppl. Fig. 9F). Interestingly, Nicheformer’s performance increased with neighborhood size in the case of the brain and liver dataset. In the liver, we observed a stronger performance trend which might be related to transcriptional patterns of zonation and structural components in the liver^90^. For the CosMx liver dataset, we additionally evaluated whether a multi-task multi-layer perceptron (MLP) would allow the prediction of all neighborhood sizes jointly (Methods). We observed that a multi-task MLP did not outperform a neighborhood size-specific linear probing model or the fine-tuned Nicheformer model, indicating that downstream tasks should be evaluated separately (Suppl. Fig. 8G).

To understand the model’s behavior and performance in more detail, we additionally assessed the fine-tuned Nicheformer performance for each cell type separately in the MERFISH mouse brain dataset (Fig. 5D, Methods). We computed the absolute error between predicted and true neighborhood compositions across all four neighborhood sizes and sorted the result based on the median values per cell type. We found that the most accurately predicted cell types in terms of absolute error are also within the eight (out of 33) most abundant cell types in the MERFISH mouse brain dataset. In contrast, the four cell types for which Nicheformer performed worse are in the 14 least abundant cell types (Fig. 5D). For example, highly abundant cell types predominantly from cortical layers (IT-ET Glut, NP-CT-L6b Glut) are structurally organized in the brain and have a quite homogeneous neighborhood composition. Those two factors help to explain the very accurate Nicheformer predictions. Similarly, CB Glut cells are based in the cerebellum, an area with very high cell density^91^ and high neighborhood homogeneity. Even though they have a lower abundance in the overall dataset, Nicheformer accurately predicted their neighborhood composition (Fig. 5D). On the other hand, Nicheformer shows a lower performance on cell types predominantly found in the midbrain or hypothalamus (MB GABA, MB, Dopa, HY Glut, Hy MM Glut). These cell types are relatively rare cell types in the given dataset and are located in more diverse and complex tissue layouts and show a greater variety of neighboring cell types^13,91^. This indicates that regionally diverse and less abundant cell types in the pretraining corpus are harder to predict for the Nicheformer model. The performance differences might be related to the structural properties of the brain regions as well as their varying cell type compositions and abundance in the dataset. We further observed a relatively good performance of Nicheformer for the neighborhood composition prediction of immune cells, despite their relatively low abundance and their lack of regional specificity in the brain. Immune cells are scattered across the brain and accomplish very specific but differing tasks ranging from regulating synaptic plasticity, and immune surveillance, to preventing excitotoxicity^92^. Interestingly, the Nicheformer embedding of the immune cells in the MERFISH mouse brain data preserves the region information of those cells and region-specific subclusters can be identified (Fig. 5E).

To assess whether our results generalize across organs and technologies, we performed a similar analysis for the CosMx human liver dataset, evaluating the overall cell type performance in the task of predicting the neighborhood composition across resolutions (Suppl. Fig. 9H). Again, we observed that Nicheformer’s performance heavily depends on the cell type abundance in the dataset and the regional specificity of the individual cells, e.g., we saw a lower absolute error for hepatocytes compared to circulating immune cells (Suppl. Fig. 9H). Hepatocytes are predominantly found in highly structured cellular microenvironments and show strong spatial patterns in their gene expression^93^. While liver-resident immune cell populations were shown to be mobilized under certain circumstances, hence their regional specificity might be lower compared to other cell types^94^. This indicates that the Nicheformer embeddings can be useful to identify and understand region- and niche-specific structures and differentiate cell types that show a higher regional specificity.

#### Nicheformer infers cellular niche density in unseen data

Beyond cellular niche labels and neighborhood composition, we asked whether local cell density is encoded in a cell’s expression profile. It is long known that cell density can strongly affect growth behavior in vivo and in culture; also, increased cell density is a key feature of the formation of the tumor microenvironment, which leads to the creation of a hypoxic environment and depletion of infiltrating immune cell populations^95,96^. For example, in colon cancer, it was shown that the immune cell density is associated with patient survival and can be used for tumor-immune patient stratification for improved anticancer therapy^87^. In non-small cell lung cancer^86^, immune cell density and neighborhood compositions were used to stratify specimens into groups associated with clinical outcomes.

We tested whether Nicheformer accurately predicts the neighborhood density in a Xenium lung dataset measuring an adult human healthy lung section and a section with invasive adenocarcinoma from a second patient^97^, and in a Xenium FFPE-preserved healthy and diseased colon with stage 2A adenocarcinoma from two different patients^97^. Consistent with literature observations^87,86^, we observed a higher average cellular density in the cancer sections (colon=12.3 cells, lung=12.1 cells) compared to healthy tissue (colon=10.7 cells, lung=10.7 cells) when extracting cellular neighborhoods at the same radius (Fig. 6A, F, Methods).

**Figure 6.**
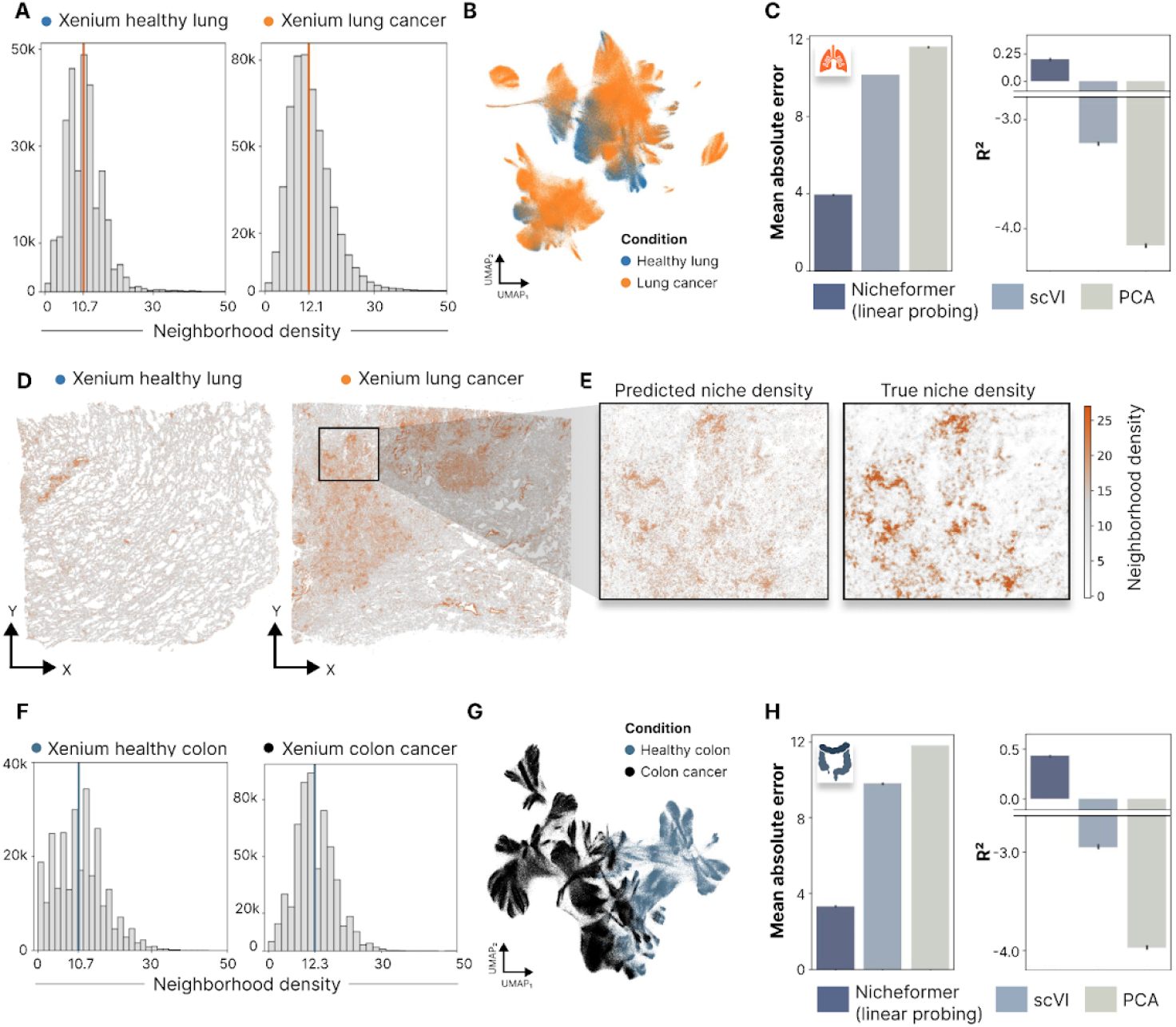
Nicheformer accurately predicts changes in cellular neighborhood density in the lung and colon. A) Barplot of cellular neighborhood densities split by condition for the Xenium human lung dataset. B) UMAP of the Nicheformer embedding of the Xenium human lung dataset colored by condition. C) Mean absolute error and R^2^ for the cellular neighborhood density prediction task for a Nicheformer linear probing model and linear probing models trained on scVI and PCA embeddings. D) Spatial allocation of cells in the Xenium human lung dataset colored by predicted cellular neighborhood density in the healthy and diseased lung. E) Predicted versus true cellular neighborhood density for a zoomed-in section of the Xenium lung cancer section. F) Barplot of cellular neighborhood densities split by condition for the Xenium human colon dataset. G) UMAP of the Nicheformer embedding of the Xenium human colon dataset colored by condition. H) Mean absolute error and R^2^ for the cellular neighborhood density prediction task for a Nicheformer linear probing model and linear probing models trained on scVI and PCA embeddings.

We first computed Nicheformer embeddings for both datasets by generating a forward pass through the Nicheformer pretrained model (Fig. 6B, G). Additionally, we embedded the two datasets with scVI, and PCA (Methods). The three resulting embeddings for the datasets are then used as input for a linear probing regression model to predict the cellular neighborhood density for each cell. The linear probing models trained on the scVI and PCA embeddings failed to correctly predict the mean density and performed worse than random prediction, resulting in negative R^2^-values for both tissues. Interestingly, the linear probing model trained on the Nicheformer embedding outperformed the other two models in terms of mean absolute error and R^2^ (Fig. 6C, H) and was able to accurately predict a higher cellular density in the tumor regions and denser tissue structures in the Xenium lung dataset (Fig. 6D). This demonstrates that the Nicheformer embeddings are able to capture neighborhood density variation solely on transcriptome information better than the baselines. Nicheformer’s ability to infer cellular neighborhood density in healthy and cancer tissue can be useful to inject spatial relationship information in dissociated data to further characterize cell state variation in systems such as the tumor microenvironment.

## Discussion

Nicheformer demonstrates the potential of multiscale foundation models for dissociated single-cell and spatial transcriptomics data. By leveraging the SpatialCorpus-110M and evaluating the model in different spatially informed downstream tasks and assessing the model’s prediction uncertainty, we demonstrate that Nicheformer captures complex relationships between gene expression and spatial context. We introduce a novel set of downstream tasks designed explicitly for spatial data analysis, in which Nicheformer consistently outperforms baseline models, including foundational models trained only on scRNA-seq data such as GeneFormer and scGPT, highlighting its effectiveness in learning a cell representation that is able to predict spatial features and the need to train on multiscale datasets to capture the intricate spatial relationships present in tissue organization. These results strongly suggest that spatial context can be effectively inferred from transcriptomics data using Nicheformer. Further, Nicheformer paves the way for transferring spatial information to large collections of dissociated single-cell data which opens the door for more nuanced analyses of cellular function in tissue environment in-silico.

A cell integrates its spatial context, i.e., its cellular neighborhood by cell communication, which is reflected in the cell’s transcriptomic profile^98^. This property has been used successfully to learn cell type communication profiles from coexpressed receptor-ligand interactions^99–101^, to reconstruct spatial gene expression from spatial context and anchor points using optimal transport^102–104^ and to determine cell interactions beyond known receptor-ligands via graph neural networks^89,105^. With Nicheformer, we build upon these results and show that we can predict spatial context from a cell’s gene expression profiles alone with consistent accuracy. We found that, for example, immune cell neighborhoods in the brain are most likely encoded in the gene expression profiles, making it easier for Nicheformer to understand these differences and relate them to neighborhood composition changes. Extending this analysis to additional tissues will enable the characterization of recurrent immune niches across tissues and organs.

A long-term vision in systems biology has been to create multiscale models, from molecules and cells up to tissue, organs and eventually the whole organism^106,107^. Nicheformer is a step towards creating a generalizable multiscale model for single-cell and spatial biology, bridging the gap from the single-cell to the tissue modality. More generally, it will be necessary to operate on multimodal data to generate a true representation of the cellular state. While spatial transcriptomics captures the cellular microenvironment in tissues well, integrating additional data modalities, such as protein abundance or epigenetic modifications, will provide a more complete picture of the cellular state. The development of multimodal foundation models faces multiple challenges. One key hurdle is the lack of sufficient paired data measured across multiple or even all cellular modalities. However, with the development of new assays and sequencing technologies, we expect the number of multimodal datasets to grow, enabling the development of architectures to model them. Incorporating additional modalities will remain a challenge in the future as e.g. epigenetic modifications, protein abundance, or gene expression all have unique characteristics, and effectively combining them in a way that leverages their strengths remains an ongoing research area.

While Nicheformer represents a milestone for learning general representations for single-cell biology, we acknowledge some of the limitations of this approach. Firstly, Nicheformer performance depends on the data abundance and transcriptional diversity of the cells under study. Indeed, we showed that Nicheformer’s performance for accurately predicting spatial labels and spatial compositions is impacted by cell and tissue type abundance in a spatial transcriptomics dataset. With the ongoing growth in spatial transcriptomics data availability as well as improved throughput thanks to technological advances, we expect that the prediction performance will increase across evaluated tissues. Secondly, Nicheformer does not explicitly incorporate the physical location of a cell during pretraining, limiting its capability to fully leverage the available information on spatial context. We deliberately chose not to include spatial coordinates during pretraining because we wanted to learn a general representation of gene expression variation across both modalities, fully supervised by gene expression alone. Nevertheless, we anticipate that future iterations of Nicheformer will account for spatial relationships of cells by encoding spatial neighbor graphs, for example, and potentially leveraging graph transformer architectures^108–110^ for the pretraining stage on spatial transcriptomics data. Graph transformers excel at modeling relationships between nodes in graphs, making them ideal for capturing nearest-neighbor effects on a cell’s transcriptome. Thirdly, the interpretability of the Nicheforme model has not been fully explored. In future iterations, it would be interesting to inspect the learned architecture in order to understand interactions between genes within cells and niches to extract biological mechanistic knowledge, for example, by assessing how gene relationships are associated with cell state across the two modalities under consideration. We additionally see a need to scale Nicheformer in the number of parameters, pretraining time and dataset size. Characterizing scaling laws for foundation models in genomics has the potential to identify bottlenecks in learning schemes and datasets, thus informing design and pretraining choices for the next generation of models. Finally, we want to highlight the need for more comprehensive benchmarks than the set of spatial tasks presented here, which will help judge extensions and future alternative models. The field of biological foundation models is a novel area brimming with potential. However, unlike more established AI domains^111,112^, there’s a crucial gap in the form of standardized benchmarks for evaluating these models. Establishing robust benchmarks is a critical next step to compare and improve performance, rigorously assess methodological progress and guide future model development to identify the full potential of foundation models for single-cell biology.

Overall, Nicheformer demonstrates the feasibility of learning a foundational representation able to effectively transfer information from single-cell to spatial genomics and its reverse, paving the way for the next generation of foundation models trained on large heterogeneous collections of dissociated and single-cell data. We describe a set of new evaluations that are explicitly designed to probe the model’s ability to encode spatial context and its transferability to a different modality that can be leveraged as a new benchmark for multimodal foundation models for single-cell and spatial genomics. We believe Nicheformer represents an important milestone towards building a general and robust representation of cellular biology phenotypes advancing our understanding of the heterogeneous effects of cellular niches in development and disease. We envision Nicheformer and similar models to actively assist in experimental design through hypothesis generation and experiment selection, ultimately accelerating the pace of scientific progress by helping to choose the next set of most informative experiments. Nicheformer will thus help to guide and design spatial experiments based on single-cell RNA-seq measurements, supporting the upcoming transition from cell to tissue atlases.

## Methods

### Collection of the SpatialCorpus-110M

#### Dissociated data collection

We collected and combined dissociated single-cell and single-nucleus data from the latest patch of CellXGene^113^, 50 additional curated studies available through the sfaira data zoo^59^, 150 datasets acquired through the Gene Expression Omnibus (GEO) data repository^57,58^ and four datasets from the Human Cell Atlas (HCA) data explorer^114^.

For the data originating from CellXGene, we used the CZ CellXGene Discover Census^113^ version 2023-07-15 and its Python API to download the latest batch of all data available on the census. The CZ CellXGene Discover Census only contains cells from human or mouse, as well as only gene expression measurements obtained via RNA sequencing. We additionally only downloaded primary data that was marked with the respective identifier in the Census to ensure that cells are not represented multiple times in our collection. Subsequently, we downloaded the entire cell and gene metadata as well as the raw counts and stored them as H5AD on disk.

For additional data acquisition, firstly, we selected human and mouse 10x Genomics technology datasets not present in the latest CellXGene patch from the sfaira data zoo, which was acquired and categorized as previously described^59^, and excluded datasets without publicly available raw count matrices. We then downloaded the selected data through the sfaira interface, removed any cells with less than 200 expressed genes, streamlined the feature space of each dataset to Ensembl release 104 (GRCh38) protein-coding genes, applied sfaira metadata streamlining, and applied the Nicheformer metadata scheme. We stored the data for each study from sfaira as individual H5AD objects on disk.

Secondly, or the acquisition from the GEO data repository, we focused on GEO IDs previously included in the recent scsimilarity^45^ preprint publication. After cross-checking this list with the other used data sources to avoid duplicated data, we acquired the necessary metadata from the GEO website and the corresponding publications. We downloaded the count matrices, converted the various data formats into AnnData format and combined them with the collected metadata to save them as individual H5AD objects on disk. We curated ontology term identifiers for species based on the ontology representation of the NCBI organismal taxonomy (NCBITaxon)^115^, tissue based on the Uber-anatomy ontology (Uberon)^116,117^, sex based on the ontology of phenotypic qualities (PATO)^118,119^ and assay based on the Experimental Factor Ontology (EFO)^120^. All ontology terms were obtained through the Ontology Lookup Service (OLS)^121^.

Lastly, we followed the same approach for the four HCA data explorer^60^ datasets as for the GEO datasets. To make the dataset acquisition process reproducible and available to the community, we have shared scripts for downloading and standardizing of all datasets. All data-collection-related code can be found at https://github.com/theislab/nicheformer-data. We additionally implemented a validator to streamline the verification process, ensuring alignment between metadata formats and the data collection schema. A detailed list and overview table of all datasets containing GEO ID, DOI, the number of cells, tissue, assay, and author information can be found in Suppl. Table 3.

#### Spatial data collection

The spatial part of the SpatialCorpus-110M consists of datasets measured with image-based spatial transcriptomics technologies, namely CosMx, ISS, MERFISH, and 10x Xenium. We collected 60 different datasets across 15 different solid organs. Most of the spatial data collection was collected via the Vizgen data release^63^, the 10x Genomics data resource^64^ and the CosMx data resource^122^. The remaining datasets were collected through the data resources stated in the original publications. Unpublished datasets were obtained prior to publication via the original authors. Each dataset was downloaded and stored as individual H5AD files. For each dataset, we collected expression data and associated gene and cell-level metadata, but high-resolution images and segmentation masks were not collected and curated. We curated ontology term identifiers for species based on the ontology representation of the NCBI organismal taxonomy (NCBITaxon)^115^, tissue based on the Uber-anatomy ontology (Uberon)^116,117^, sex based on the ontology of phenotypic qualities (PATO)^118,119^ and assay based on the Experimental Factor Ontology (EFO)^120^. All ontology terms were obtained through the Ontology Lookup Service (OLS)^121^. For Xenium and CosMx assays official ontology terms are not yet defined, so we replaced them with placeholders. For datasets that did not provide Ensembl gene identifiers, we used pyEnsembl^65^ with the Ensembl release 104 (GRCh38) to map gene names to Ensembl gene identifiers and subsequently BioMart^66^ through the official Ensembl releases^67^ for mapping mouse genes to orthologous gene identifiers. Scripts for acquiring the spatial data are also shared in our GitHub repository. We used the same validator as used for the dissociated datasets to streamline the verification process of the collected metadata. We applied no additional quality control, gene or cell level filtering for the spatial omics datasets beyond the filters applied by the original authors of the publications or the filters automatically applied by the individual spatial transcriptomics technologies. A detailed list and overview table containing the GEO ID, DOI, the number of cells, tissue, assay, and author information for the spatial datasets can be found in Suppl. Table 4.

#### Datasets used for downstream tasks and evaluations

Publically available datasets used for downstream tasks and evaluations were collected in the same way as the other spatial transcriptomics datasets present in the SpatialCorpus-110M. As most of our downstream tasks require cell type, niche and region label annotations, we focussed primarily on annotated and large-scale spatial transcriptomics datasets. We provide a detailed description of those datasets below.

### MERFISH mouse brain

Yao et al.^13^ measured 4.3 million cells across 59 tissue sections from one whole male mouse brain using multiplexed error-robust fluorescence in situ hybridization (MERFISH) with a 500-gene panel. This dataset contains a hierarchical cell type annotation structured into four nested levels of annotation. We used the *class_label* field with 33 distinct cell types as input for the Nicheformer niche regression task (Suppl. Fig. 3C), the *division_id* label, containing seven distinct labels (CBX-MOB-other neuronal, Immune, Low Quality - LQ, Neuroglial, PAL-sAMY-TH-HY-MB-HB neuronal, Pallium glutamatergic, Subpallium GABAergic, Vascular), as niche labels (Suppl. Fig. 7B), and the *clean_region_label* field, containing 17 distinct labels (CB, CTXsp, HB, HIP, HY, Isocortex, LSX, MB, OLF, PAL, RHP, STRd, STRv, TH, sAMY, ventricle, white_matter), as the region label (Suppl. Fig. 7A) for the Nicheformer label prediction tasks. The tissue niches represent the cellular organization in the brain, grouping together neurons by major brain structure (pallium, subpallium, hypothalamus/extended amygdala, thalamus/epiphysis, and mid/hindbrain) as well as major neurotransmitter type (glutamate and GABA)^13^. Non-neuronal cells are grouped into neuroglial, immune, and vascular niches. The train-test split defined for this dataset is composed of a random image or tissue section hold-out across all sections in the measured entire male mouse brain (Suppl. Fig. 7A-C).

### CosMx human liver

We collected the CosMx human liver dataset from the publicly available CosMx data resource^122^. The dataset comprises cells from both a normal healthy liver measuring 332,877 cells across 301 fields of views covering one tissue section in a male 35-year-old patient, as well as cells from a Hepatocellular Carcinoma measuring 460,441 cells across 383 fields of view in one tissue section from a 65-year-old female patient. Both samples were measured with the 1000-plex CosMx Human Universal Cell Characterization Panel. The dataset includes both cell type and niche labels. For the niche label prediction task, we used the healthy liver section, which provides six distinct labels defining structural zones in the liver: Portal vein (Zone 1a), Zone 1b, Zone 2a, Zone 2b, Zone 3a, and Central vein (Zone 3b) (Suppl. Fig. 9B, D). We did not use the cancer liver sample for the niche label prediction task as it was primarily composed of cells annotated as a general tumor niche without further substructures provided. For the niche composition prediction task, we used both the cancer and healthy liver sections with the cell type labels, which define 22 distinct cell types (Antibody secreting B cells, CD3+ alpha beta T cells, Central venous LSECs, Cholangiocytes, Erthyroid cells, Hep, Hep 1, Hep 3, Hep 4, Hep 5, Hep 6, Inflammatory macrophages, Mature B cells, NK like cells, Non inflammatory macrophages, Periportal LSECs, Portal endothelial cells, Stellate cells, gamma delta T cells 1, tumor 1, tumor 2 and an undefined type (NotDet) (Suppl. Fig. 9E). The train-test split defined for this dataset is composed of a random field of view hold-out across both tissue sections (Suppl. Fig. 9A, D).

### CosMx human lung

We collected the CosMx human lung dataset from the publicly available CosMx data resource^122^. This dataset contains samples from five different donors (301,611, 89,975, 227,110, 71,304 and 81,236 cells, respectively) across eight fields of views measured with the 1000-plex CosMx Human Universal Cell Characterization Panel. All donors have just one field of view, with the exception of the first donor, which has three fields of view and the third, which has two fields of view. The train-test split defined for this dataset is composed of a random field of view hold-out (Suppl. Fig. 11A-B). CosMx provides both cell type and niche labels. We use the 22 distinct cell type labels defined in this dataset for the niche composition prediction task. These labels are B-cell, NK, T CD4 memory, T CD4 naive, T CD8 memory, T CD8 naive, Treg, endothelial, epithelial, fibroblast, mDC, macrophage, mast, monocyte, neutrophil, pDC, plasmablast, tumor 12, tumor 13, tumor 5, tumor 6, tumor 9 (Suppl. Fig. 11C).

### Xenium human lung

We collected the Xenium human lung dataset from the 10x Genomics data resource (https://www.10xgenomics.com/datasets). This dataset measures two different lung sections, an adult human healthy lung (295,883 cells) and an adult human lung with invasive adenocarcinoma (531,165 cells). Both sections are measured with the 289-plex Xenium Human Lung Gene Expression Panel and an additional 100 lung-cell type-specific genes. As this dataset is not annotated, we only use it for the neighborhood density prediction task. We computed a spatial graph of cells with a radius of 25 µm^2^ to calculate the cellular niche densities. The train-test split defined for this dataset is a random cell hold-out across all cells from both sections.

### Xenium human colon

We collected the Xenium human colon dataset from the 10x Genomics data resource (https://www.10xgenomics.com/datasets). This dataset measures two different colon FFPE-preserved tissue sections: a non-diseased colon (275,822 cells) and a cancer stage 2A adenocarcinoma (587,115 cells). Both sections are measured with the 325-plex Xenium Human Colon Gene Expression Panel and an additional 100 genes specifically selected to cover signaling and chemokine genes, and markers for stromal cells. As again this dataset is not annotated, we only use it for the neighborhood density prediction task. We computed a spatial graph of cells with a radius of 17 µm^2^ in both sections to calculate the cellular niche densities. The train-test split defined for this dataset is a random cell hold-out across all cells from both sections.

### Dissociated dataset used for label transfer

#### Single-cell RNA-seq of the primary motor cortex

Yao et al. generated a large-scale transcriptomic and epigenetic atlas of the mouse primary motor cortex^14^. We subsetted this large-scale dataset to cells measured with 10x v3 single-cell RNA seq. The subset captures 21,884 genes in 7,416 cells and annotates 19 different cell types (Astro, Endo, L5 ET, L5 IT, L6 CT, L6 IT, L6 IT Car3, L6b, L2/3 IT, L5/6 NP, Lamp5, Microglia, OPC, Oligo, Pvalb, Sncg, Sst, CLMC and Vip) (Fig. 3C). We manually transferred cell types present in this dataset to the cell types measured in the MERFISH mouse brain dataset. Respectively, we mapped Astro to Astro-Epen; Endo and VLMC to Vascular; Microglia to Immune; Oligo and OPC to Oligo; L6 IT, L6 IT *Car3*, L5 IT, L2/3 IT, L5 ET to IT-ET Glut; L5/6 NP, L6b and L6 CT to NP-CT-L6b Glut; and Lamp5, Sncg, Vip Pvalb and Sst to CGE/MGE GABA.

### Nicheformer tokenization, architecture and pretraining

#### Nicheformer tokenization

The Nicheformer training corpus encompasses over 110 million cells in total, measured in more than 350 datasets using 8 different sequencing technologies and two species: human and mouse. The total number of genes considered is 20,310, being 16,981 orthologous genes, 3,178 human-specific and 151 mouse-specific. For Nicheformer, we use a tokenization strategy similar to the one in Geneformer^2^ with the difference that the cell transcripts are normalized according to the technology-specific non-zero mean to account for differences in the sequencing protocol. First, all cells are normalized so that each of them has 10,000 counts. To account for technological variations, we then compute a technology-specific gene expression non-zero mean vector i.e., the mean expression value of each gene, without considering the zero counts. We computed a single dissociated mean expression vector for the dissociated datasets because the differences between sequencing protocols in the dissociated cells are not as large as in the spatial assays. We then normalize the expression of each cell using the corresponding technology-specific mean expression vector to obtain the expression of each gene in each cell relative to the whole training corpus. Finally, the genes are ranked in descending order, from most expressed to lowest expressed, excluding all not expressed genes, creating an ordered set *T* of genes:

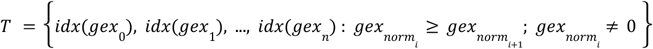

where *idx*(*gex*_*i*_) is a function that returns the index of gene *i* in a previously defined vocabulary of genes and *gex*_*i*_ is the gene expression of gene *i* of a cell. To incorporate the influence of biological context on gene expression, we prepend contextual tokens for <ASSAY>, <MODALITY> and <ORGANISM> to the set *T* to incorporate metadata information to the input data. These tokens encode metadata information, such as assay type (e.g., MERFISH, CosMx, 10x 5’ v2, etc.), modality (dissociated or spatial), and organism (mouse or human). Recognizing the significant impact biological context can have on gene expression, we augment the input sequences for our transformer model with modality, organism, and assay tokens. This approach allows the model to explicitly learn representations that account for context-driven variations, leading to more robust and generalizable downstream analyses. Therefore, for a cell *i*, with a specific assay, organism and modality, the ordered set of tokens *T*^*i*^ is the following:

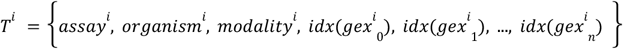

As a last step, the length of the set *T*^*i*^ is truncated to *N* = 1500. As not all cells have the same number of expressed genes, there might be sets whose total length is lower than 1500. In those cases, <PAD> tokens are appended such that the final length is *N* = 1500. <PAD> tokens ensure that all inputs have the same length by filling empty spaces with no semantic meaning. This is an important element when handling cells belonging to both RNA-seq and spatial assays because gene panels are usually smaller in the latter, which leads to a larger amount of <PAD> tokens in the set.

#### Nicheformer architecture

Given an initial input set 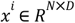 composed of *N* tokens of dimensionality *D*, Nicheformer encodes the position within the set by adding positional embeddings. Instead of modeling as sinusoidal embeddings, we use learnable embeddings for each position^19^.

Nicheformer is composed of 12 stacked transformer blocks such that the output of one block is in the input of the following block. Given an input sequence 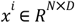:

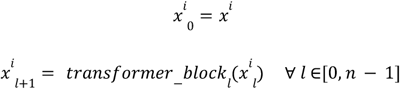

Each transformer block consists of two main modules: a multihead self-attention mechanism and a feed-forward neural network. The multi-head self-attention mechanism enables the model to weigh the relevance of different input elements in the input set when generating output representations^27^. In our case, we use 16 attention heads, token dimensionality *D*=512 and dimensionality of the hidden layer of the FFN of 1024. The <PAD> tokens are masked for the attention mechanism so that no token can pay attention to them.

#### Nicheformer pretraining and performance optimization

Nicheformer optimizes masked language modeling (MLM) loss^19^ during pretraining. We mask 15% of the tokens, including contextual and gene tokens but excluding <PAD> tokens, during pretraining. The model is then trained to predict the original tokens that have been masked, utilizing the unmasked tokens as context. Specifically, following the BERT schema^19^, if the i-th token is chosen to be masked, 80% of the time it is replaced by a <MASK> token, 10% of the time by another random gene or contextual token and 10% of the time it remains unchanged. Mathematically, the MLM loss is described as:

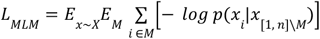

where *M* is the set of masked tokens, *X* is the entire dataset, *x* is a cell of the dataset and *x*_*i*_ is gene *i* of the cell *x*.

Nicheformer was pretrained for approximately 10 days using three compute nodes, each with four Nvidia A100 40GB GPUs (total 12 GPUs). We train the model using bfloat16 mixed precision. We use the AdamW optimizer^123^ with β_1_ = 0. 9 and β_2_ = 0. 999, weight decay of 0.1, and dropout of 0.0. The batch size is nine and the gradients are accumulated during 10 batches before running the backward pass. The minimum learning rate is 1e-5, which increases until 1e-3 with a linear warmup of 100,000 steps. After the warmup, a cosine decay regime^124^ is applied. Gradient clipping is set to 1.0 during the first epoch and then decreased to 0.5. All weights are initialized using Xavier initialization^125^ with default parameters, while the bias terms are initialized to 0. Checkpoints were taken every 10000 steps.

### Downstream tasks

#### Spatial cell type, niche and region label prediction

For the spatial cell type, niche and region label classification task, we use the respective labels defined in the individual datasets (see section on Datasets used for downstream tasks and evaluations). We extracted the unique labels for each class, transferred them to 64-bit signed integer values and one-hot encoded them as a matrix with n different classes, with n being either the number of cell types, niches or region. We then used for linear probing a linear layer optimized with a cross-entropy loss. We trained on the training set of the respective dataset for one epoch at a learning rate of 1e-3 and with a batch size of 256. The performance metrics reported are calculated on a held-out test set. We selected the model-assigned class label by calculating the argmax over the output vector of the linear layer. Classification uncertainties reported in this work are the output of the linear layer rescaled to [0,1] such that the sum equals 1 using a Softmax function.

#### Neighborhood composition

For the neighborhood composition regression tasks, we first define a spatial graph of cells by building an adjacency matrix based on the Euclidean distance in the two-dimensional coordinate space provided by the individual datasets. The adjacency matrix of spatial cells is a block-diagonal matrix *A ∈ R*^*nxn*^, with *n* the number of cells present in the dataset calculated based on the spatial proximity of cells where connectivities can only occur within a field of view. We use a binary adjacency matrix with *a*_*ij*_ = 1 if *d*(*x*_*i*_, *x* _*j*_) ≤ δ_*r*_ where *d*(*·, ·*) describes the Euclidean distance between nodes *i, j ∈ n* and δ_*r*_ the is the maximal distance between cells, and *a*_*ij*_ = 0 otherwise. We do not include self-connectivities for the adjacency matrix to not confound the signal. We additionally define the matrix of observed cell types *X*_*l*_ *∈* {0, 1}^*nxl*^ as a one-hot encoding of the *l* distinct cell types present in the dataset. The neighborhood composition for a given radius is then given as

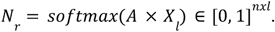

The resulting matrix reflects for each cell captured in the dataset a vector giving the proportions of cell types present in the neighborhood of the cell. For the neighborhood prediction task we used a for linear probing a linear layer followed by a Softmax function to rescale the prediction to lie in the range [0,1] and sum to 1. We used the mean square error loss for optimizing this linear layer, trained on the training set of the respective dataset for one epoch at a learning rate of 1e-3 and with a batch size of 256. The performance metrics reported are calculated on a held-out test set.

#### Neighborhood cell density prediction

For the cellular niche density, we again use the adjacency matrix of spatial cells *A ∈ R*^*nxn*^ calculated based on the Euclidean distance in the two-dimensional coordinate space. The cellular neighborhood density is then simply given by the row-wise sum of all connectivities in the adjacency matrix.

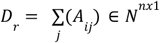

for all cells present in the dataset with *r* a given radius, *i* the index cell for which we want to calculate the density, and *j* the total number of potential neighboring cells present in the dataset. For the density prediction task, we used for linear probing a linear layer with input being the respective embedding of a cell (Nicheformer, scVI or PCA) and output a scalar. We used the mean square error loss for optimizing this linear layer, trained on the training set of the respective dataset for one epoch at a learning rate of 1e-3 and with a batch size of 256. The performance metrics reported are calculated on a held-out test set.

### Nicheformer evaluation, linear probing and fine-tuning

Nicheformer can be fine-tuned or used for linear probing. In both settings, we only train on the previously defined training set of the respective datasets used for downstream tasks (see section on Datasets used for downstream tasks and evaluation). We use in both scenarios all Nicheformer gene tokens extracted from the last layer and average them to get a cell representation. Importantly, the contextual tokens are not used in the aggregation. In linear probing, the previously computed parameter weights of the Nicheformer pretraining model are frozen, i.e., not updated further, and are subsequently used as input to a downstream task. The cell’s representation is then fed into a linear layer specific to each downstream task, which represents either a classification task in the case of the niche and region label prediction or a regression for predicting the neighborhood composition and cellular density. For the neighborhood composition task, we additionally fitted a multi-task multilayer perceptron (MLP) that uses the Nicheformer embedding as input and predicts the varying neighborhood composition vectors in a dataset. The MLP is optimized using the average mean squared error across all neighborhood sizes considered. Fine-tuning generally describes using a pretrained model, and training it to a specific downstream task of choice. We speak of a fine-tuned Nicheformer version when we allow the model to change the previously learned parameter space and the weights are updated for a specific task. Importantly, each downstream task can also be optimized with respect to a new set of metrics. All runs are trained for a single epoch with a maximum learning rate of 1e-4 and a cosine decay scheduler reaching 1e-5 at the end. Batch size is nine with gradients accumulated for 10 batches (Suppl. Table 5). We highlight the respective tasks and metrics used to compute them in the section Downstream task.

### Nicheformer cell embedding stability analysis

We conduct an analysis to study Nicheformer’s cell representation robustness to perturbations in the input data, that might correspond to incomplete gene panels or measurement noises. To do so, we tokenize one dissociated brain dataset and one spatial brain dataset. We then apply perturbations to the tokenized cells before feeding them into Nicheformer to obtain embeddings. Specifically, we either randomly shuffle 10%, 20%, 50%, or 100% of the genes in the gene ranking sequence (Suppl. Fig. 1A), or we drop out 10%, 20%, 50%, or 80% of the genes from the sequence (Suppl. Fig. 1B). To evaluate whether the perturbed cell embeddings still cluster with the original (non-perturbed) embeddings, we use the silhouette score from scIB^32^, using the cell type annotations provided by the authors to define the clusters.

### Nicheformer attention analysis

We conduct an attention analysis to explore how Nicheformer differentiates between male and female cells by focusing on sex-specific gene variations. For this, we feed 2000 male cells and 2000 female cells into the model and extract attention matrices from all 16 attention heads across the 12 transformer blocks. Our analysis then centers on two gene sets: a prior-knowledge set of SDGs, known for exhibiting sex differences, and a randomly sampled control set of 97 genes. Then, to assess general trends in attention distribution, we average the attention scores to obtain an attention score per layer. In addition to this, we extract the maximum attention value for each gene per layer, isolating the highest level of focus from any single attention head. Evaluating both average and maximum attention, allows us to discern whether certain genes consistently receive attention across multiple heads or are sharply focused on by individual heads. Specifically, we compare the attention scores:

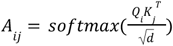

where *A*_*ij*_ represents the attention that token *i* pays to token *j*. As we have 16 attention layers, we denote A_*ij*_^*h*^ the attention that token *i* pays to token *j* in the layer *h*.

In Nicheformer, with 12 layers, the attention matrices for each layer and head are represented as *A*_*ij*_ ^(*l*,*h*)^, where *l∈*{1, 2,…, 12} represents the layer, and *h∈*{1, 2,…, 16} denotes the head. To assess how much attention each tokens pay to a token *m*, we focus on extracting the attention scores *A*_*im*_ ^(*l*,*h*)^, which capture the attention that each token *i* allocates to the *m* in layer *l* and head *h*.

For each observation, we compute both the maximum and average attention that any token *i* pays to the token *m* across all heads in each layer. This is done by first calculating the maximum and average attention for each layer as follows:

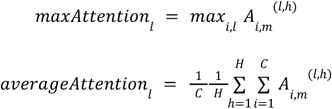

where *I* refers to all other tokens in the sequence and *H* is the number of heads (16). These values give us, respectively, the highest attention score and the average attention score that the token *m* receives from other tokens for each layer, considering all heads. By averaging these maximum and average attention values across multiple observations, we can assess how attention is distributed across layers, identifying the layers where the token *m* receives the most focus and how consistently it receives attention across tokens and heads.

### Benchmarking against competing methods

#### Comparisons against Geneformer and scGPT

To get the Geneformer embeddings we used the release version 0.0.1 of the official Geneformer repository on HuggingFace and extracted the embeddings using the pre-trained weights of the larger 12 layer variant provided at the time. We used the second to last layer to get a more general representation as recommended by the repository. We also used mean pooling as the only available option provided to aggregate the output gene embeddings into a single cell embedding.

For the comparison against scGPT, we first created scGPT embeddings using scGPT 0.2.1, pre-trained on the whole human as recommended in the original publication. The embeddings were generated for three datasets, the MERFISH mouse brain, the CosMx human lung and the CosMx human liver. For the MERFISH mouse dataset we first mapped the mouse genes to human genes using BioMart^66^ through the official Ensembl releases^67^. The fraction of overlapping genes compared to the gene context used in scGPT was for the MERFISH mouse brain data 471/483 genes, for the CosMx human liver dataset 997/999 genes, and for the CosMx human lung dataset 958/960 genes.

The resulting Geneformer and scGPT embeddings then served as input to a linear layer specific to each downstream task (Suppl. Table 5).

#### Baseline comparisons to scVI and PCA embeddings

We compared the performance of the fine-tuned Nicheformer model and the linear probing scenario to embeddings generated with scVI^29^ and PCA. We generated scVI and PCA embeddings on just the downstream datasets themselves and additionally on an informed 1% subset of all datasets present in the SpatialCorpus-110M. We used this subset to train two different scVI models as specified in (Suppl. Table 5) to generate latent representations with 512 and 10 dimensions respectively. The two models were then used to obtain latent representations for the datasets that were used for downstream task evaluations. The PCA embeddings were generated in a similar way using the implementation available in sklearn version 1.4.1 to obtain PCA embeddings of dimensions 512 and 10 respectively.

We split the fine-tuning datasets (MERFISH mouse brain, CosMx human liver, CosMx human lung, Xenium human lung, Xenium human colon) into a training and test set, using the same random splits as applied for the Nicheformer fine-tuning. scVI and PCA were computed on each fine-tuning dataset individually. We used scvi-tools version 1.1.2 with a negative binomial distribution gene likelihood on the raw gene expression counts and trained scVI on the training set with a batch size of 256 for 10 epochs and used 2 hidden layers for the encoder and decoder neural networks. The resulting embedding was chosen to have a latent dimension of 256. After training, we returned the latent representation for each cell in both the training and the test set.

For generating principal component analysis (PCA) embeddings for each dataset we used the implementation available in sklearn version 1.4.1. We first normalized the respective raw gene expression counts for each dataset so that each cell has a total number of counts equal to the median of the total counts for all cells with scanpy version 1.10.1. Next we use scanpy to log1p-transform the data matrix to ensure the data is centered before using it as input to the PCA implementation. We use the sklearn implementation with the number of components being set to 30. All other parameters are the defaults provided by the sklearn implementation. We fit the PCA on the training set and afterwards apply the dimensionality reduction to both the training and test set. The resulting lower dimensional representations, X_scvi and X_pca then serve as input to a linear layer specific to each downstream task (Suppl. Table 5).

## Code and data availability

All models described here are implemented in a Python package available at https://github.com/theislab/nicheformer. Downloading and preprocessing scripts for all public datasets used in pretraining and fine-tuning the models will be available at https://github.com/theislab/nicheformer-data.

The Allen brain atlas consortium generated the Allen Institute brain atlas mouse p20, Allen Institute brain atlas mouse p28, and Allen Institute brain atlas mouse female datasets (Suppl. Table 4), which have kindly been provided to us prior to publication. The Xenium human spinal cord, the ISS human brain GBM, the ISS human discover healthy lung, the ISS mouse EAE MS, and the Xenium mouse brain datasets have been generated by Mat Nillson lab and have been kindly provided to us prior to publication.

## Author contributions

F.J.T. conceived the study with the help of A.C.S., and A.T.L.; A.C.S. and A.T.L. contributed equally and have the right to list their name first in their curriculum vitae; F.J.T. supervised the project; A.T.L. and A.C.S. performed the analysis and wrote the code; A.T.L. led the data engineering, model design, implementation and pretraining in discussion with G.P., L.H. and A.C.S. led the data curation effects for the dissociated data collection; A.C.S. led the data curation efforts for the spatial data collection; M.M. and L.D. supported the data curation efforts for the dissociated data collection; F.D. and R.G. supported the data curation efforts for the spatial data collection; R.G. and L.V. helped to interpret the brain and liver results; A.C.S. designed and created all main figures; A.C.S., A.T.L., G.P., R.G., and F.J.T. wrote the manuscript. All authors read, corrected, and approved the manuscript.

## Acknowledgments

We thank Luke Zappia, Leon Hetzel, Alessandro Palma, Sara Jimenéz, Felix Fischer, Jan Engelmann, Artur Szałata, Lukas Heumos, and Merel Kuijs for valuable discussions and feedback on this project. We thank lamin.ai, specifically Alex Wolf, Lukas Heumos, Sunny Sun, and Sergei Rybakov for helpful discussions on data curation, data management and model training. Additionally, we would like to thank Hongkui Zeng, Michael Kunst, and the Allen Brain Atlas consortium for providing us early access to their MERFISH whole mouse brain atlas as well as the additional unpublished MERFISH mouse brain datasets. Additionally, we would like to thank Mats Nilsson and Sergio Marco Salas for providing us early access to their unpublished Xenium and ISS datasets.

This work was co-funded by the European Union (ERC, DeepCell - 101054957) and supported by the Chan Zuckerberg Initiative Foundation (CZIF; grant CZIF2022-007488 (Human Cell Atlas Data Ecosystem)), by the Wellcome Leap ΔTissue Program and through the BRAIN Initiative Cell Atlas Network (BICAN). G.P. and

L.D. acknowledge funding by the Joachim Herz Foundation. L.D. was additionally supported by the BMBF-funded de.NBI Cloud within the German Network for Bioinformatics Infrastructure (de.NBI) (031A532B, 031A533A, 031A533B, 031A534A, 031A535A, 031A537A, 031A537B, 031A537C, 031A537D, 031A538A).

## References

1. Cui, H. et al. scGPT: toward building a foundation model for single-cell multi-omics using generative AI. Nat. Methods (2024) doi:10.1038/s41592-024-02201-0.

2. Theodoris, C. V. et al. Transfer learning enables predictions in network biology. Nature 618, 616–624 (2023).

3. Lopez, R., Regier, J., Cole, M. B., Jordan, M. I. & Yosef, N. Deep generative modeling for single-cell transcriptomics. Nat. Methods 15, 1053–1058 (2018).

4. Sikkema, L. et al. An integrated cell atlas of the lung in health and disease. Nat. Med. 29, 1563–1577 (2023).

5. Kanemaru, K. et al. Spatially resolved multiomics of human cardiac niches. Nature 619, 801–810 (2023).

6. Zhang, P. & Li, S. Human cross-tissue cell atlases: unprecedented resources towards systematic understanding of physiology and diseases. Signal Transduct Target Ther 7, 352 (2022).

7. Domínguez Conde, C. et al. Cross-tissue immune cell analysis reveals tissue-specific features in humans. Science 376, eabl5197 (2022).

8. Williams, C. G., Lee, H. J., Asatsuma, T., Vento-Tormo, R. & Haque, A. An introduction to spatial transcriptomics for biomedical research. Genome Med. 14, 68 (2022).

9. Du, J. et al. Advances in spatial transcriptomics and related data analysis strategies. J. Transl. Med. 21, 330 (2023).

10. Marx, V. Method of the Year: spatially resolved transcriptomics. Nat. Methods 18, 9–14 (2021).

11. Fischer, D. S., Schaar, A. C. & Theis, F. J. Modeling intercellular communication in tissues using spatial graphs of cells. Nat. Biotechnol. (2022) doi:10.1038/s41587-022-01467-z.

12. Varrone, M., Tavernari, D., Santamaria-Martínez, A., Walsh, L. A. & Ciriello, G. CellCharter reveals spatial cell niches associated with tissue remodeling and cell plasticity. Nat. Genet. 56, 74–84 (2024).

13. Yao, Z. et al. A high-resolution transcriptomic and spatial atlas of cell types in the whole mouse brain. Nature 624, 317–332 (2023).

14. Yao, Z. et al. A transcriptomic and epigenomic cell atlas of the mouse primary motor cortex. Nature 598, 103–110 (2021).

15. He, S. et al. High-plex imaging of RNA and proteins at subcellular resolution in fixed tissue by spatial molecular imaging. Nat. Biotechnol. 40, 1794–1806 (2022).

16. Lu, Y. et al. Spatial transcriptome profiling by MERFISH reveals fetal liver hematopoietic stem cell niche architecture. Cell Discov 7, 47 (2021).

17. Bommasani, R. et al. On the opportunities and risks of foundation models. arXiv preprint 2108.07258 (2021).

18. Brown, T. B. et al. Language Models are Few-Shot Learners. arXiv [cs.CL] (2020) doi:10.48550/ARXIV.2005.14165.

19. Devlin, J., Chang, M.-W., Lee, K. & Toutanova, K. BERT: Pre-training of Deep Bidirectional Transformers for Language Understanding. arXiv [cs.CL] (2018).

20. Dosovitskiy, A. et al. An Image is Worth 16×16 Words: Transformers for Image Recognition at Scale. arXiv [cs.CV] (2020) doi:10.48550/ARXIV.2010.11929.

21. Gemini Team et al. Gemini: A Family of Highly Capable Multimodal Models. arXiv [cs.CL] (2023).

22. Ji, Y., Zhou, Z., Liu, H. & Davuluri, R. V. DNABERT: pre-trained Bidirectional Encoder Representations from Transformers model for DNA-language in genome. Bioinformatics 37, 2112–2120 (2021).

23. Yang, F. et al. scBERT as a large-scale pretrained deep language model for cell type annotation of single-cell RNA-seq data. Nature Machine Intelligence 4, 852–866 (2022).

24. Rives, A. et al. Biological structure and function emerge from scaling unsupervised learning to 250 million protein sequences. Proc. Natl. Acad. Sci. U. S. A. 118, (2021).

25. Lin, Z. et al. Evolutionary-scale prediction of atomic-level protein structure with a language model. Science 379, 1123–1130 (2023).

26. Madani, A. et al. Large language models generate functional protein sequences across diverse families. Nat. Biotechnol. 41, 1099–1106 (2023).

27. Vaswani, A. et al. Attention Is All You Need. arXiv [cs.CL] (2017).

28. Ericsson, L., Gouk, H., Loy, C. C. & Hospedales, T. M. Self-Supervised Representation Learning: Introduction, Advances and Challenges. arXiv [cs.LG] (2021).

29. Gayoso, A. et al. A Python library for probabilistic analysis of single-cell omics data. Nat. Biotechnol. 40, 163–166 (2022).

30. Virshup, I. et al. The scverse project provides a computational ecosystem for single-cell omics data analysis. Nat. Biotechnol. (2023) doi:10.1038/s41587-023-01733-8.

31. Eraslan, G., Simon, L. M., Mircea, M., Mueller, N. S. & Theis, F. J. Single-cell RNA-seq denoising using a deep count autoencoder. Nat. Commun. 10, 390 (2019).

32. Luecken, M. D. et al. Benchmarking atlas-level data integration in single-cell genomics. Nat. Methods 19, 41–50 (2022).

33. Lotfollahi, M. et al. Mapping single-cell data to reference atlases by transfer learning. Nat. Biotechnol. 40, 121–130 (2022).

34. Roohani, Y., Huang, K. & Leskovec, J. Predicting transcriptional outcomes of novel multigene perturbations with GEARS. Nat. Biotechnol. (2023) doi:10.1038/s41587-023-01905-6.

35. Lotfollahi, M., Wolf, F. A. & Theis, F. J. scGen predicts single-cell perturbation responses. Nat. Methods 16, 715–721 (2019).

36. Lotfollahi, M. et al. Predicting cellular responses to complex perturbations in high-throughput screens. Mol. Syst. Biol. 19, e11517 (2023).

37. Hetzel, L. et al. Predicting Cellular Responses to Novel Drug Perturbations at a Single-Cell Resolution. arXiv [cs.LG] (2022).

38. Cui, H., Wang, C., Maan, H., Duan, N. & Wang, B. scFormer: A Universal Representation Learning Approach for Single-Cell Data Using Transformers. bioRxiv 2022.11.20.517285 (2022) doi:10.1101/2022.11.20.517285.

39. Rosen, Y. et al. Universal cell embeddings: A foundation model for cell biology. bioRxiv (2023) doi:10.1101/2023.11.28.568918.

40. Chen, J. et al. Transformer for one stop interpretable cell type annotation. Nat. Commun. 14, 223 (2023).

41. Wen, H. et al. CellPLM: Pre-training of Cell Language Model Beyond Single Cells. bioRxiv 2023.10.03.560734 (2023) doi:10.1101/2023.10.03.560734.

42. Hou, W. & Ji, Z. Assessing GPT-4 for cell type annotation in single-cell RNA-seq analysis. bioRxiv (2023) doi:10.1101/2023.04.16.537094.

43. Hao, M. et al. Large Scale Foundation Model on Single-cell Transcriptomics. bioRxiv 2023.05.29.542705 (2023) doi:10.1101/2023.05.29.542705.

44. Consens, M. E. et al. To Transformers and Beyond: Large Language Models for the Genome. arXiv preprint 2311.07621 (2023).

45. Heimberg, G. et al. Scalable querying of human cell atlases via a foundational model reveals commonalities across fibrosis-associated macrophages. bioRxiv (2023) doi:10.1101/2023.07.18.549537.

46. He, F. et al. Parameter-Efficient Fine-Tuning Enhances Adaptation of Single Cell Large Language Model for Cell Type Identification. bioRxiv (2024) doi:10.1101/2024.01.27.577455.

47. Khan, S. A. et al. Reusability report: Learning the transcriptional grammar in single-cell RNA-sequencing data using transformers. Nature Machine Intelligence 5, 1437–1446 (2023).

48. Liu, T., Li, K., Wang, Y., Li, H. & Zhao, H. Evaluating the Utilities of Foundation Models in Single-cell Data Analysis. bioRxiv (2024) doi:10.1101/2023.09.08.555192.

49. Boiarsky, R., Singh, N., Buendia, A., Getz, G. & Sontag, D. A Deep Dive into Single-Cell RNA Sequencing Foundation Models. bioRxiv 2023.10.19.563100 (2023) doi:10.1101/2023.10.19.563100.

50. Kedzierska, K. Z., Crawford, L., Amini, A. P. & Lu, A. X. Assessing the limits of zero-shot foundation models in single-cell biology. bioRxiv 2023.10.16.561085 (2023) doi:10.1101/2023.10.16.561085.

51. Alsabbagh, A. R. et al. Foundation Models Meet Imbalanced Single-Cell Data When Learning Cell Type Annotations. bioRxiv 2023.10.24.563625 (2023) doi:10.1101/2023.10.24.563625.

52. Yang, X. et al. GeneCompass: deciphering universal gene regulatory mechanisms with a knowledge-informed cross-species foundation model. Cell Res. 1–16 (2024).

53. Cook, D. P. et al. A Comparative Analysis of Imaging-Based Spatial Transcriptomics Platforms. bioRxiv 2023.12.13.571385 (2023) doi:10.1101/2023.12.13.571385.

54. Hartman, A. & Satija, R. Comparative analysis of multiplexed in situ gene expression profiling technologies. bioRxiv (2024) doi:10.1101/2024.01.11.575135.

55. Lopez, R. et al. A joint model of unpaired data from scRNA-seq and spatial transcriptomics for imputing missing gene expression measurements. arXiv preprint 1905.02269 (2019).

56. Salas, S. M. et al. Optimizing Xenium In Situ data utility by quality assessment and best practice analysis workflows. bioRxiv 2023.02.13.528102 (2023) doi:10.1101/2023.02.13.528102.

57. Edgar, R., Domrachev, M. & Lash, A. E. Gene Expression Omnibus: NCBI gene expression and hybridization array data repository. Nucleic Acids Res. 30, 207–210 (2002).

58. Barrett, T. et al. NCBI GEO: archive for functional genomics data sets--update. Nucleic Acids Res. 41, D991–5 (2013).

59. Fischer, D. S. et al. Sfaira accelerates data and model reuse in single cell genomics. Genome Biol. 22, 248 (2021).

60. HCA Data Explorer. https://explore.data.humancellatlas.org/.

61. Chen, K. H., Boettiger, A. N., Moffitt, J. R., Wang, S. & Zhuang, X. RNA imaging. Spatially resolved, highly multiplexed RNA profiling in single cells. Science 348, aaa6090 (2015).

62. Ke, R. et al. In situ sequencing for RNA analysis in preserved tissue and cells. Nat. Methods 10, 857–860 (2013).

63. Data release program. Vizgen https://vizgen.com/data-release-program/ (2021).

64. Datasets. 10x Genomics https://www.10xgenomics.com/datasets.

65. Perkins, A. & Henze, C. Increasing the efficiency of GEOS-Chem Adjoint model runs using a Python ensemble manager. (2012).

66. Smedley, D. et al. BioMart--biological queries made easy. BMC Genomics 10, 1–12 (2009).

67. Martin, F. J. et al. Ensembl 2023. Nucleic Acids Res. 51, D933–D941 (2023).

68. Tsukahara, S. & Morishita, M. Sexually Dimorphic Formation of the Preoptic Area and the Bed Nucleus of the Stria Terminalis by Neuroestrogens. Front. Neurosci. 14, 545195 (2020).

69. Guerra-Cantera, S. et al. Sex Differences in Metabolic Recuperation After Weight Loss in High Fat Diet-Induced Obese Mice. Front. Endocrinol. 12, 796661 (2021).

70. Ngun, T. C., Ghahramani, N., Sánchez, F. J., Bocklandt, S. & Vilain, E. The genetics of sex differences in brain and behavior. Front. Neuroendocrinol. 32, 227–246 (2011).

71. Brager, D. H., Sickel, M. J. & McCarthy, M. M. Developmental sex differences in calbindin-D(28K) and calretinin immunoreactivity in the neonatal rat hypothalamus. J. Neurobiol. 42, (2000).

72. Immenschuh, J. et al. Sex differences in distribution and identity of aromatase gene expressing cells in the young adult rat brain. Biol. Sex Differ. 14, 54 (2023).

73. Morrison, H. W. & Filosa, J. A. Sex differences in astrocyte and microglia responses immediately following middle cerebral artery occlusion in adult mice. Neuroscience 339, 85–99 (2016).

74. Yagi, S. et al. Sex Differences in Maturation and Attrition of Adult Neurogenesis in the Hippocampus. eNeuro 7, (2020).

75. Liu, X., Porteous, R. & Herbison, A. E. Robust GABAergic Regulation of the GnRH Neuron Distal Dendron. Endocrinology 164, bqac194 (2022).

76. Wang, L. & Moenter, S. M. Differential Roles of Hypothalamic AVPV and Arcuate Kisspeptin Neurons in Estradiol Feedback Regulation of Female Reproduction. Neuroendocrinology 110, 172–184 (2020).

77. Arendt, D. et al. The origin and evolution of cell types. Nat. Rev. Genet. 17, 744–757 (2016).

78. Heumos, L. et al. Best practices for single-cell analysis across modalities. Nat. Rev. Genet. 1–23 (2023).

79. Palla, G., Fischer, D. S., Regev, A. & Theis, F. J. Spatial components of molecular tissue biology. Nat. Biotechnol. 40, 308–318 (2022).

80. Satija, R., Farrell, J. A., Gennert, D., Schier, A. F. & Regev, A. Spatial reconstruction of single-cell gene expression data. Nat. Biotechnol. 33, 495–502 (2015).

81. Pham, D. et al. Robust mapping of spatiotemporal trajectories and cell-cell interactions in healthy and diseased tissues. Nat. Commun. 14, 7739 (2023).

82. Hu, J. et al. SpaGCN: Integrating gene expression, spatial location and histology to identify spatial domains and spatially variable genes by graph convolutional network. Nat. Methods 18, 1342–1351 (2021).

83. Mages, S. et al. TACCO unifies annotation transfer and decomposition of cell identities for single-cell and spatial omics. Nat. Biotechnol. 41, 1465–1473 (2023).

84. Wang, Q. et al. The Allen Mouse Brain Common Coordinate Framework: A 3D Reference Atlas. Cell 181, 936–953.e20 (2020).

85. Feichtenbeiner, A. et al. Critical role of spatial interaction between CD8^+^ and Foxp3^+^ cells in human gastric cancer: the distance matters. Cancer Immunol. Immunother. 63, 111–119 (2014).

86. Barua, S. et al. Spatial interaction of tumor cells and regulatory T cells correlates with survival in non-small cell lung cancer. Lung Cancer 117, 73–79 (2018).

87. Galon, J. et al. Type, density, and location of immune cells within human colorectal tumors predict clinical outcome. Science 313, 1960–1964 (2006).

88. Fridman, W. H., Pagès, F., Sautès-Fridman, C. & Galon, J. The immune contexture in human tumours: impact on clinical outcome. Nat. Rev. Cancer 12, 298–306 (2012).

89. Fischer, D. S., Schaar, A. C. & Theis, F. J. Learning cell communication from spatial graphs of cells. bioRxiv 2021.07.11.451750 (2021) doi:10.1101/2021.07.11.451750.

90. Hildebrandt, F. et al. Spatial Transcriptomics to define transcriptional patterns of zonation and structural components in the mouse liver. Nat. Commun. 12, 7046 (2021).

91. Zhang, M. et al. Molecularly defined and spatially resolved cell atlas of the whole mouse brain. Nature 624, 343–354 (2023).

92. Colonna, M. & Butovsky, O. Microglia Function in the Central Nervous System During Health and Neurodegeneration. Annu. Rev. Immunol. 35, 441–468 (2017).

93. Ben-Moshe, S. & Itzkovitz, S. Spatial heterogeneity in the mammalian liver. Nat. Rev. Gastroenterol. Hepatol. 16, 395–410 (2019).

94. Robinson, M. W., Harmon, C. & O’Farrelly, C. Liver immunology and its role in inflammation and homeostasis. Cell. Mol. Immunol. 13, 267–276 (2016).

95. The evolving tumor microenvironment: From cancer initiation to metastatic outgrowth. Cancer Cell 41, 374–403 (2023).

96. Parra, E. R. et al. Immune cellular patterns of distribution affect outcomes of patients with non-small cell lung cancer. Nat. Commun. 14, 2364 (2023).

97. [No title]. https://www.10xgenomics.com/datasets/xenium-human-lung-preview-data-1-standard.

98. Armingol, E., Officer, A., Harismendy, O. & Lewis, N. E. Deciphering cell–cell interactions and communication from gene expression. Nat. Rev. Genet. 22, 71–88 (2020).

99. Efremova, M., Vento-Tormo, M., Teichmann, S. A. & Vento-Tormo, R. CellPhoneDB: inferring cell-cell communication from combined expression of multi-subunit ligand-receptor complexes. Nat. Protoc. 15, 1484–1506 (2020).

100. Cang, Z. & Nie, Q. Inferring spatial and signaling relationships between cells from single cell transcriptomic data. Nat. Commun. 11, 2084 (2020).

101. Dimitrov, D. et al. LIANA+: an all-in-one cell-cell communication framework. bioRxiv 2023.08.19.553863 (2023) doi:10.1101/2023.08.19.553863.

102. Nitzan, M., Karaiskos, N., Friedman, N. & Rajewsky, N. Gene expression cartography. Nature 576, 132–137 (2019).

103. Klein, D. et al. Mapping cells through time and space with moscot. bioRxiv 2023.05.11.540374 (2023) doi:10.1101/2023.05.11.540374.

104. Haviv, D., Gatie, M., Hadjantonakis, A.-K., Nawy, T. & Pe’er, D. The covariance environment defines cellular niches for spatial inference. bioRxiv (2023) doi:10.1101/2023.04.18.537375.

105. Tanevski, J., Flores, R. O. R., Gabor, A., Schapiro, D. & Saez-Rodriguez, J. Explainable multi-view framework for dissecting inter-cellular signaling from highly multiplexed spatial data. bioRxiv 2020.05.08.084145 (2020) doi:10.1101/2020.05.08.084145.

106. Vandereyken, K., Sifrim, A., Thienpont, B. & Voet, T. Methods and applications for single-cell and spatial multi-omics. Nat. Rev. Genet. 24, 494–515 (2023).

107. Rood, J. E., Maartens, A., Hupalowska, A., Teichmann, S. A. & Regev, A. Impact of the Human Cell Atlas on medicine. Nat. Med. 28, 2486–2496 (2022).

108. Yun, S., Jeong, M., Kim, R., Kang, J. & Kim, H. J. Graph Transformer Networks. in Advances in Neural Information Processing Systems (eds. Wallach, H. et al.) vol. 32 (Curran Associates, Inc., 2019).

109. Zhao, Q., Ren, W., Li, T., Xu, X. & Liu, H. GraphGPT: Graph Learning with Generative Pre-trained Transformers. arXiv preprint 2401.00529 (2023).

110. Liu, J. et al. Towards graph foundation models: A survey and beyond. arXiv preprint 2310.11829 (2023).

111. Srivastava, A. et al. Beyond the Imitation Game: Quantifying and extrapolating the capabilities of language models. arXiv [cs.CL] (2022).

112. Chiang, W.-L. et al. Chatbot Arena: An Open Platform for Evaluating LLMs by Human Preference. arXiv [cs.AI] (2024).

113. CZI Single-Cell Biology Program et al. CZ CELL×GENE Discover: A single-cell data platform for scalable exploration, analysis and modeling of aggregated data. bioRxiv 2023.10.30.563174 (2023) doi:10.1101/2023.10.30.563174.

114. Projects - HCA Data Explorer. https://explore.data.humancellatlas.org/projects.

115. Federhen, S. The NCBI Taxonomy database. Nucleic Acids Res. 40, D136–43 (2012).

116. Mungall, C. J., Torniai, C., Gkoutos, G. V., Lewis, S. E. & Haendel, M. A. Uberon, an integrative multi-species anatomy ontology. Genome Biol. 13, R5 (2012).

117. Haendel, M. A. et al. Unification of multi-species vertebrate anatomy ontologies for comparative biology in Uberon. J. Biomed. Semantics 5, 21 (2014).

118. Gkoutos, G. V., Schofield, P. N. & Hoehndorf, R. The anatomy of phenotype ontologies: principles, properties and applications. Brief. Bioinform. 19, 1008–1021 (2018).

119. Gkoutos, G. V., Green, E. C. J., Mallon, A.-M., Hancock, J. M. & Davidson, D. Using ontologies to describe mouse phenotypes. Genome Biol. 6, R8 (2005).

120. Malone, J. et al. Modeling sample variables with an Experimental Factor Ontology. Bioinformatics 26, 1112–1118 (2010).

121. Ontology lookup service (OLS). https://www.ebi.ac.uk/ols4/.

122. He, S. et al. High-plex Multiomic Analysis in FFPE at Subcellular Level by Spatial Molecular Imaging. bioRxiv 2021.11.03.467020 (2022) doi:10.1101/2021.11.03.467020.

123. Loshchilov, I. & Hutter, F. Decoupled Weight Decay Regularization. arXiv [cs.LG] (2017).

124. Loshchilov, I. & Hutter, F. Sgdr: Stochastic gradient descent with warm restarts. arXiv preprint 1608.03983 (2016).

125. Glorot, X. & Bengio, Y. Understanding the difficulty of training deep feedforward neural networks. In Proceedings of the Thirteenth International Conference on Artificial Intelligence and Statistics (eds. Teh, Y.W. & Titterington, M.) vol. 9 249–256 (PMLR, Chia Laguna Resort, Sardinia, Italy, 13--15 May 2010).

