## Supplementary Material for "Nicheformer: a foundation model for single-cell and spatial omics"

### Supplements

#### Supplementary Figures

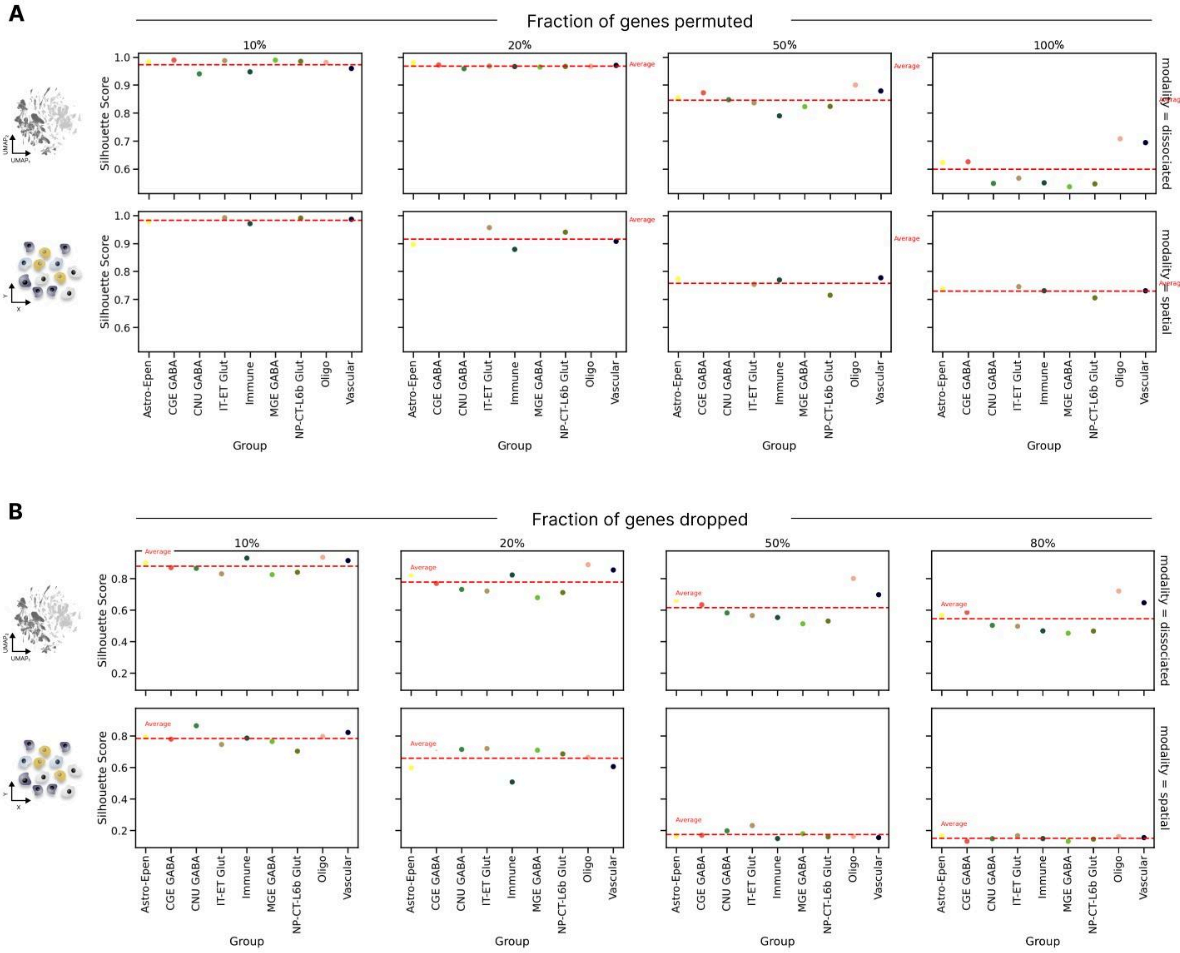


**Suppl. Figure 1 | Nicheformer's cell representations are robust to input noise. A)** We compute Nicheformer cell representations for a dissociated and spatial brain dataset and use author cell type annotations as ground truth. We randomly permute 10%, 20%, 50% and 100% of the genes in the input sequence and obtain cell representations. Then, we compute the silhouette score to evaluate how perturbed are the cell representations. **B)** We repeat the same experiment but instead of permuting genes, we drop them off the input sequence (which contains only non-zero genes). In particular, we drop 10%, 20%, 50% and 80% of the genes in the input sequence. In this case, the deterioration of the cell embeddings happens faster than when permuting genes. Cell representations of spatial cells deteriorate faster than dissociated cells (<0.2 silhouette score against >0.6 silhouette score for 50% dropout level). We hypothesise that this happens due to the shorter gene panels, i.e. in large gene panels, Nicheformer can leverage more information from the longer context length to correct disturbances in the data.

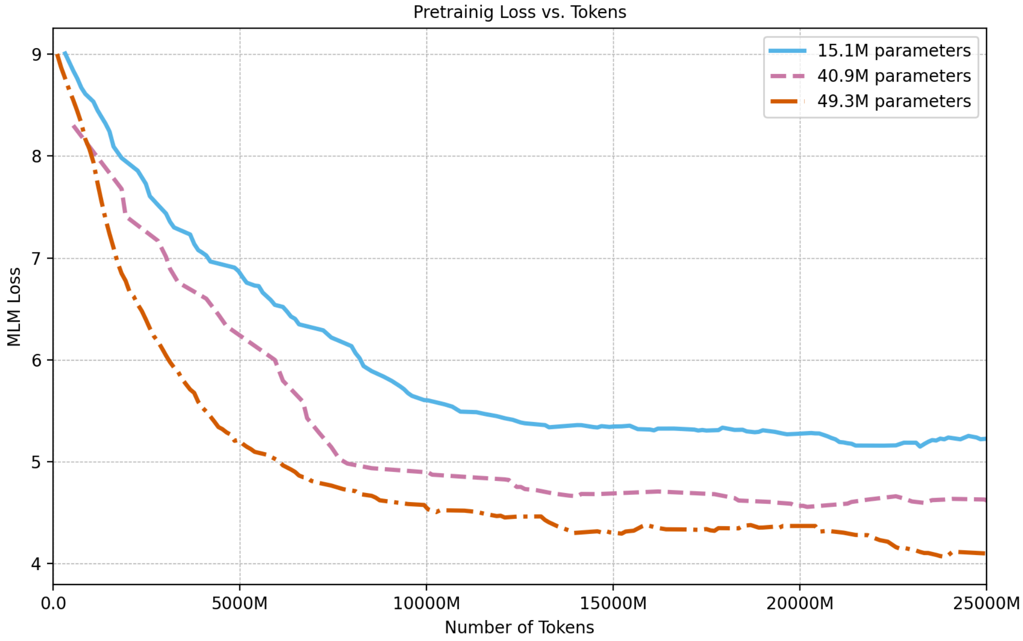


**Suppl. Figure 2 | MLM loss as a function of the total number of tokens seen by the model.** Shown are the loss curves of three different models with varying parameter size, 15.1 million parameters, 40.9 million parameters and 49.3 million parameters, respectively. The larger the model, the lower is the pretraining loss. All the losses are a moving average with a window of 10. All the models were evaluated in the same training set with fixed random seed.

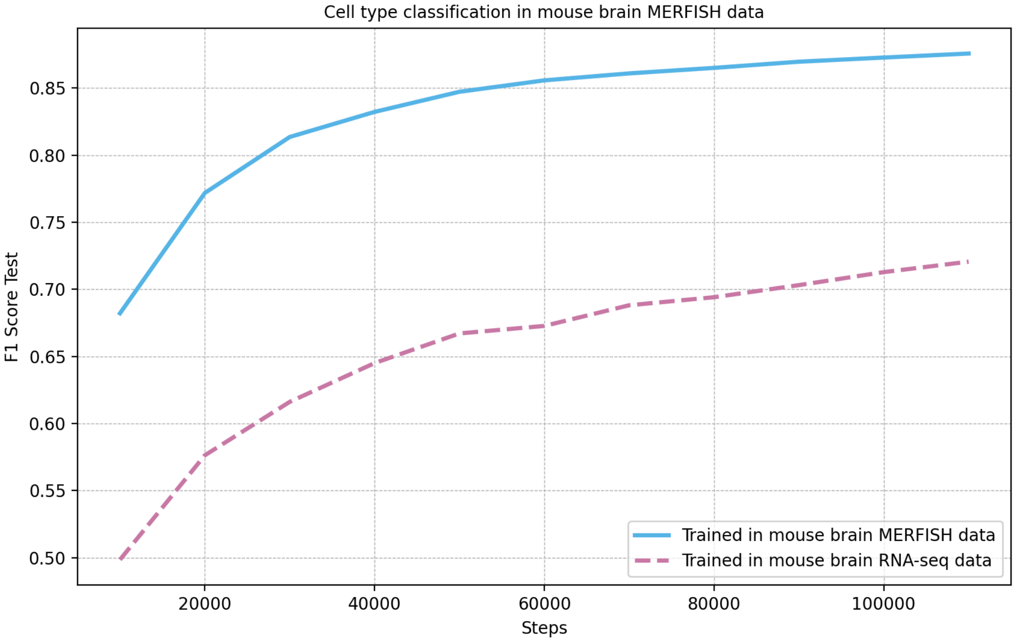


**Suppl. Figure 3 | F1 Score for cell type classification in a mouse brain MERFISH dataset.** Shown are the F1 score curves of two different models trained on different modalities: spatial and dissociated respectively. Both models have the same number of parameters and have been training for the same amount of time. The task is performed by linear probing. The model trained on MERFISH data notably outperforms the model trained on RNA-seq, highlighting a significant distribution shift between technologies.

**A**

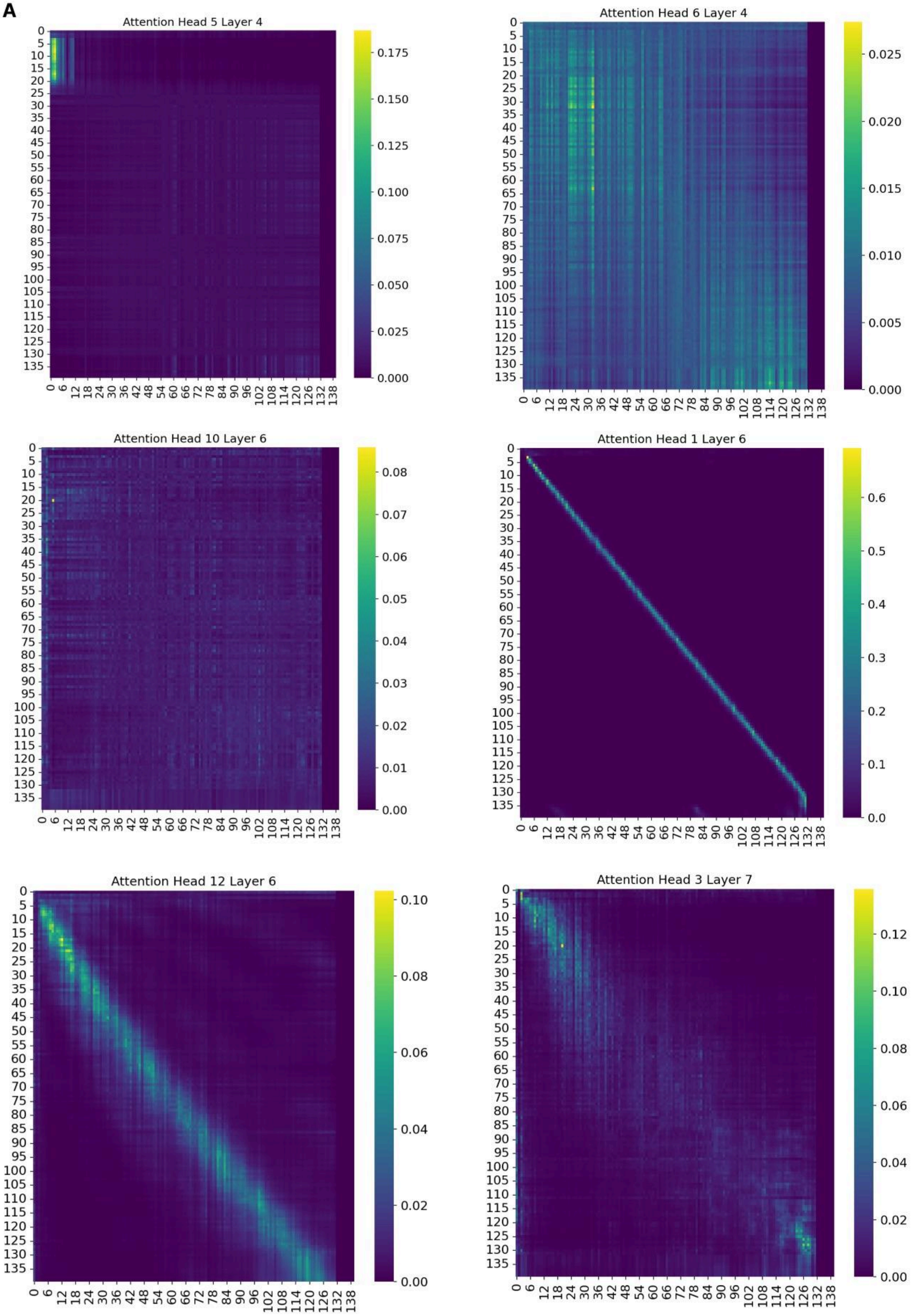


**Suppl. Figure 4| Attention matrices of different Nicheformer attention heads.** Shown are different attention matrices obtained when feeding Nicheformer with cells from the AVPV section. Different heads showcase different patterns, which reveal diverse attention behaviours, including metadata token focus (Head 5, Layer 4), selective gene interactions (Head 6, Layer 4), diffuse attention across genes (Head 10, Layer 6), strong self-attention (Head 1, Layer 6), combined self and global attention (Head 12, Layer 6), and concentrated attention on key genes (Head 3, Layer 7).

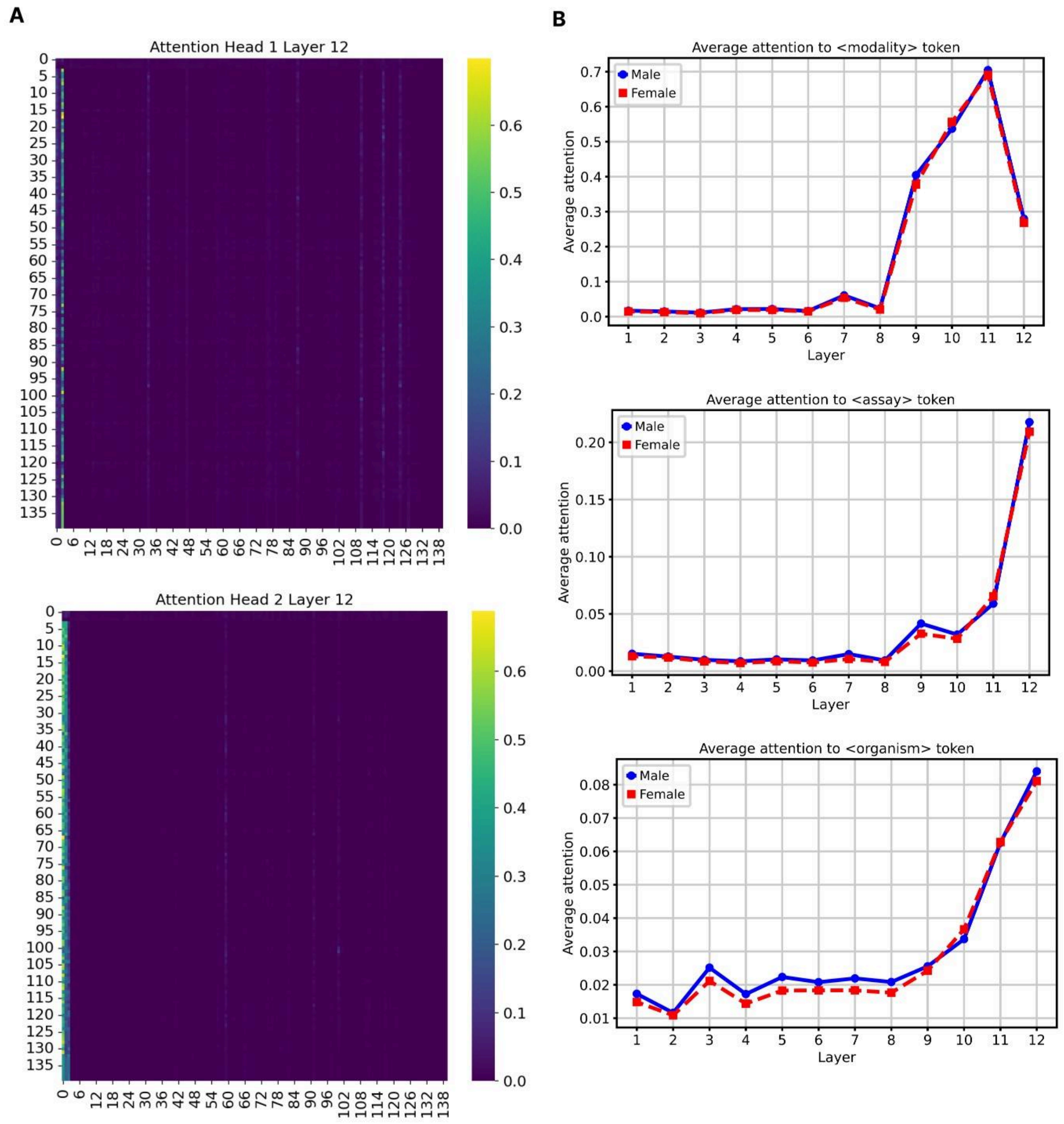


**Suppl. Figure 5| The metadata tokens are specially attended in the last transformer blocks. A)** Shown are different attention matrices extracted from the last transformer block of Nicheformer. They present a similar pattern in which almost all attention is paid to the metadata tokens. **B)** Average attention paid, per layer, to the metadata tokens. It can be observed a clear trend: the last layers of the model pay, by a large margin, the most attention to the metadata tokens. The analysis is done in both male and female brain mouse datasets to showcase that the pattern is consistent.

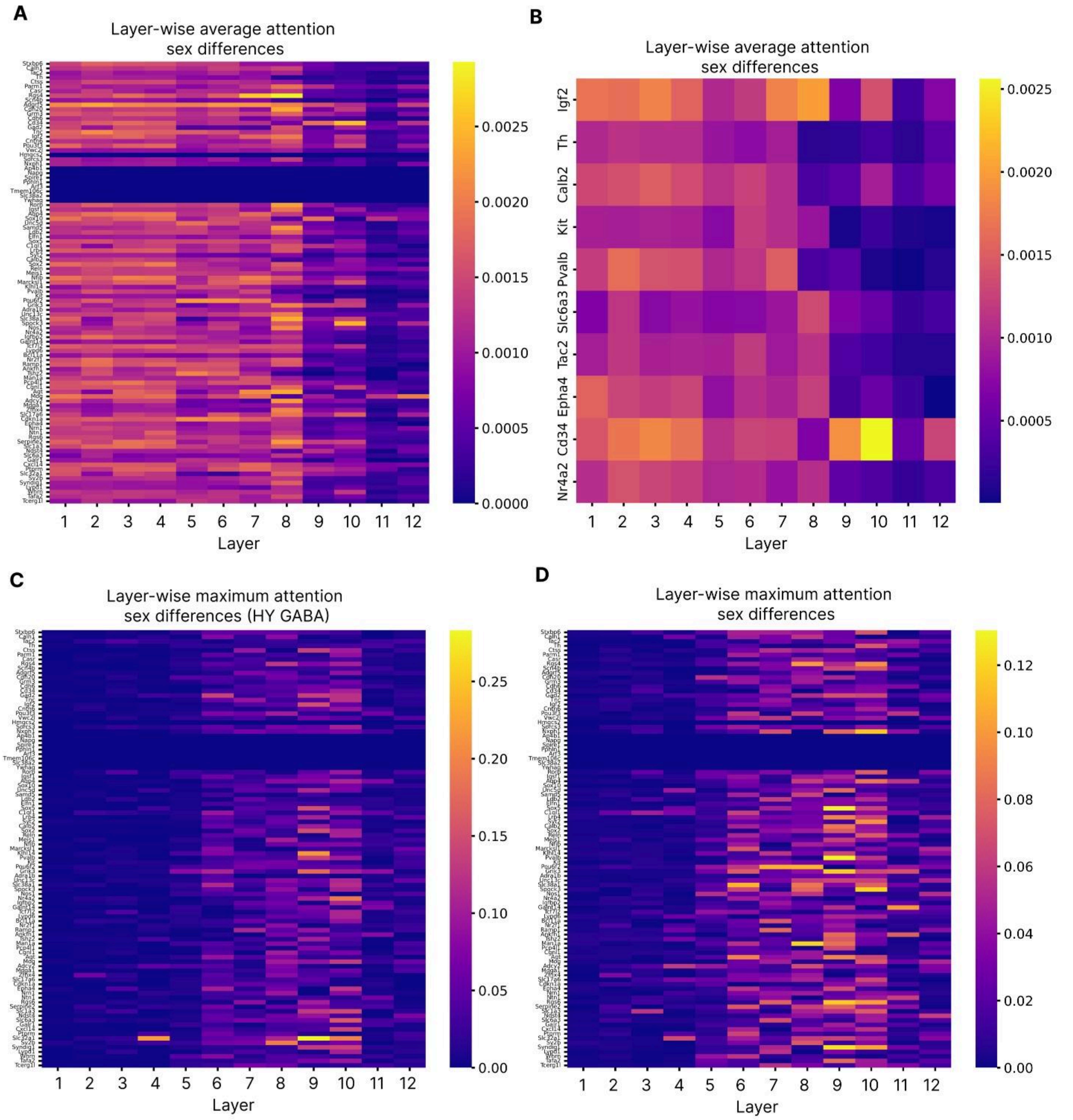


**Suppl. Figure 6| Layer-wise average attention difference between cell and female AVPV cells. A)** The first layers of Nicheformer show the highest attention differences between cell and female cells, even though this is very small. **B)** The same pattern holds for the SDN genes. **C)** Nicheformer's middle layers show the maximum attention score differences between the male and the female cells for the HY GABA cells within the AVPV section. **D)** The same pattern occurs when examining the maximum differences for all cells in the AVPV section. The contrast of the average attention difference plotted here and the maximum attention differences (Fig 3.D-F) suggests that the sex differences are captured by a subset of the attention heads. The average attention difference is computed averaging all attention heads, whereas the maximum attention difference attends to the maximum difference reported in any head.

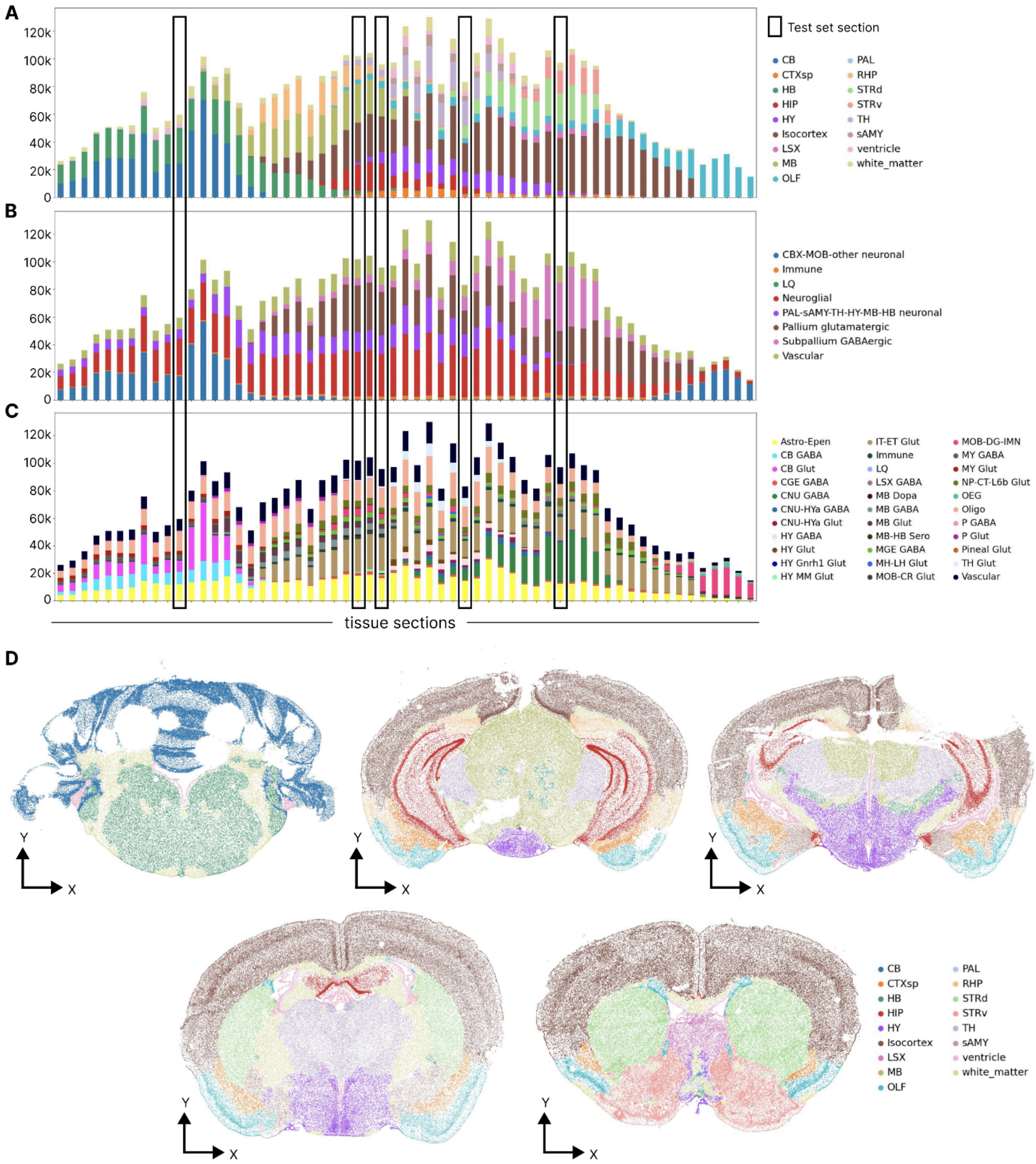


**Suppl. Figure 7 | Nicheformer fine-tuning datasets - MERFISH mouse brain. A-C)** Region (A), niche (B), and cell type (C) label distribution across all tissue sections in the MERFISH mouse brain data with the test set highlighted. D) Spatial allocation of cells in the five test tissue sections of the MERFISH mouse brian

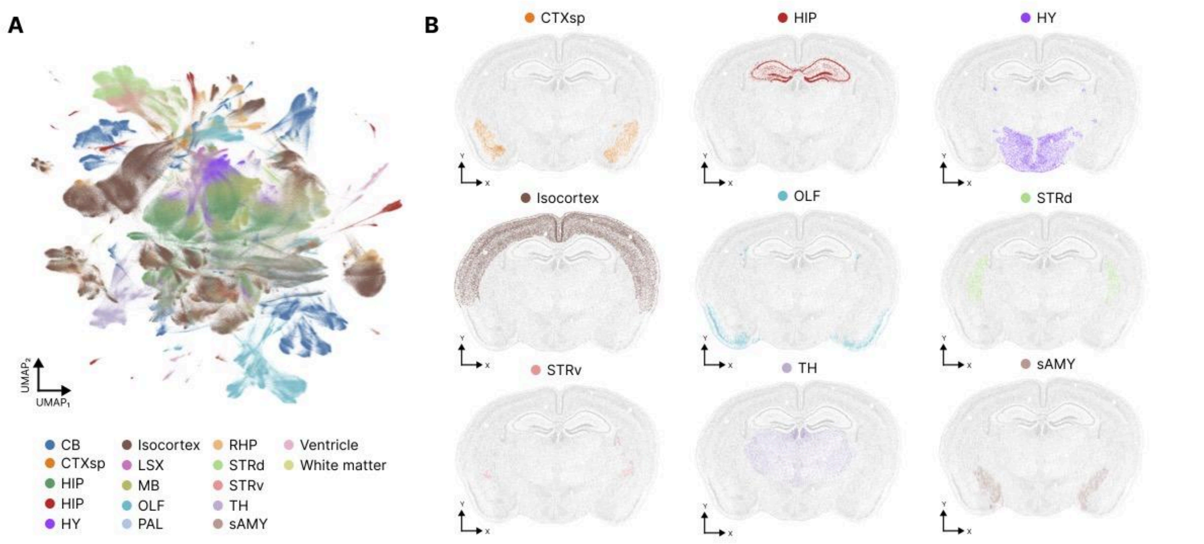


**Suppl. Figure 8 | Nicheformer embedding of the MERFISH mouse brain dataset. A)** UMAP visualization of the Nicheformer embedding of the MERFISH mouse brain dataset colored by region label. B) Exemplary brain slice of the MERFISH mouse brain dataset colored by region label.

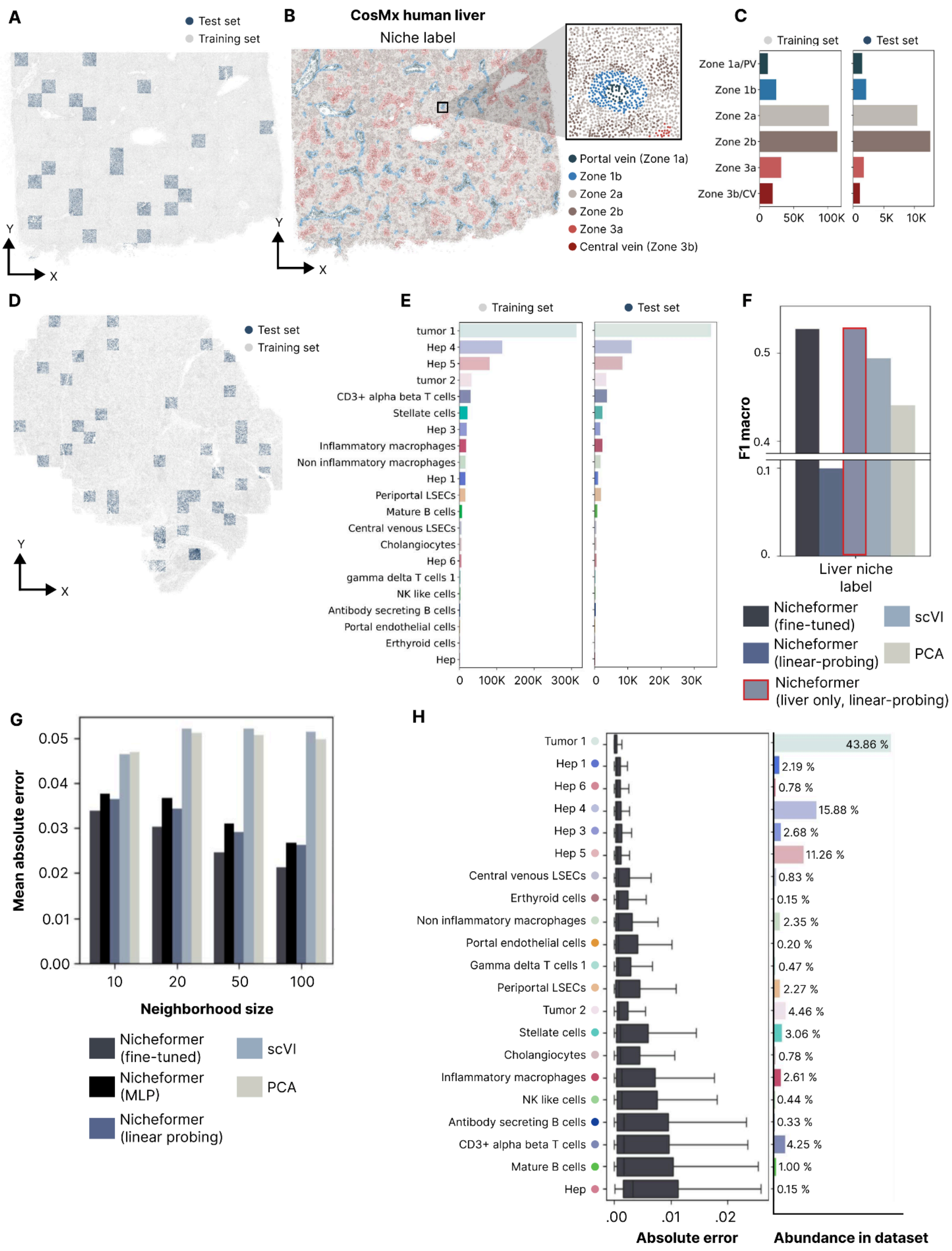


**Suppl. Figure 9 | Nicheformer fine-tuning datasets - CosMx human liver.** A-B) Spatial allocation of cells in the healthy CosMx liver section colored by training and test split used for training Nicheformer (A) and niche label (B). C) Niche label distribution in the training and test set for the healthy CosMx liver dataset. D) Spatial allocation of cells in the cancer CosMx liver section colored by training and test split used for training Nicheformer. and the cancer CosMx liver section (bottom). D) Distribution of cell type labels in the healthy and cancer CosMx liver data in both training and test set. E) Test-set F1-macro of niche label prediction of the fine-tuned Nicheformer model, the linear probing model, the linear probing model evaluated on a Nicheformer model longer trained in the liver training-set, and a linear probing baseline computed based on embeddings generated with scVI and PCA, respectively. G) The fine-tuned, a multi-task MLP on top of the Nicheformer embedding and the linear probing Nicheformer models outperform zero-shot models trained on scVI and PCA embeddings in terms of mean absolute error across all neighborhood sizes and all three organs, the brain, liver, and lung. H) Left: Fine-tuned Nicheformer performance on the CosMx human liver data grouped by index cell type. Shown are the absolute error values between predicted and observed niche composition vectors for held-out test cells. For each box in (G), the centerline defines the median, the height of the box is given by the interquartile range (IQR), the whiskers are given by $1.5 \times$ IQR and outliers are given as points beyond the minimum or maximum whisker. Right: Index cell type abundances in the entire CosMx human liver dataset.

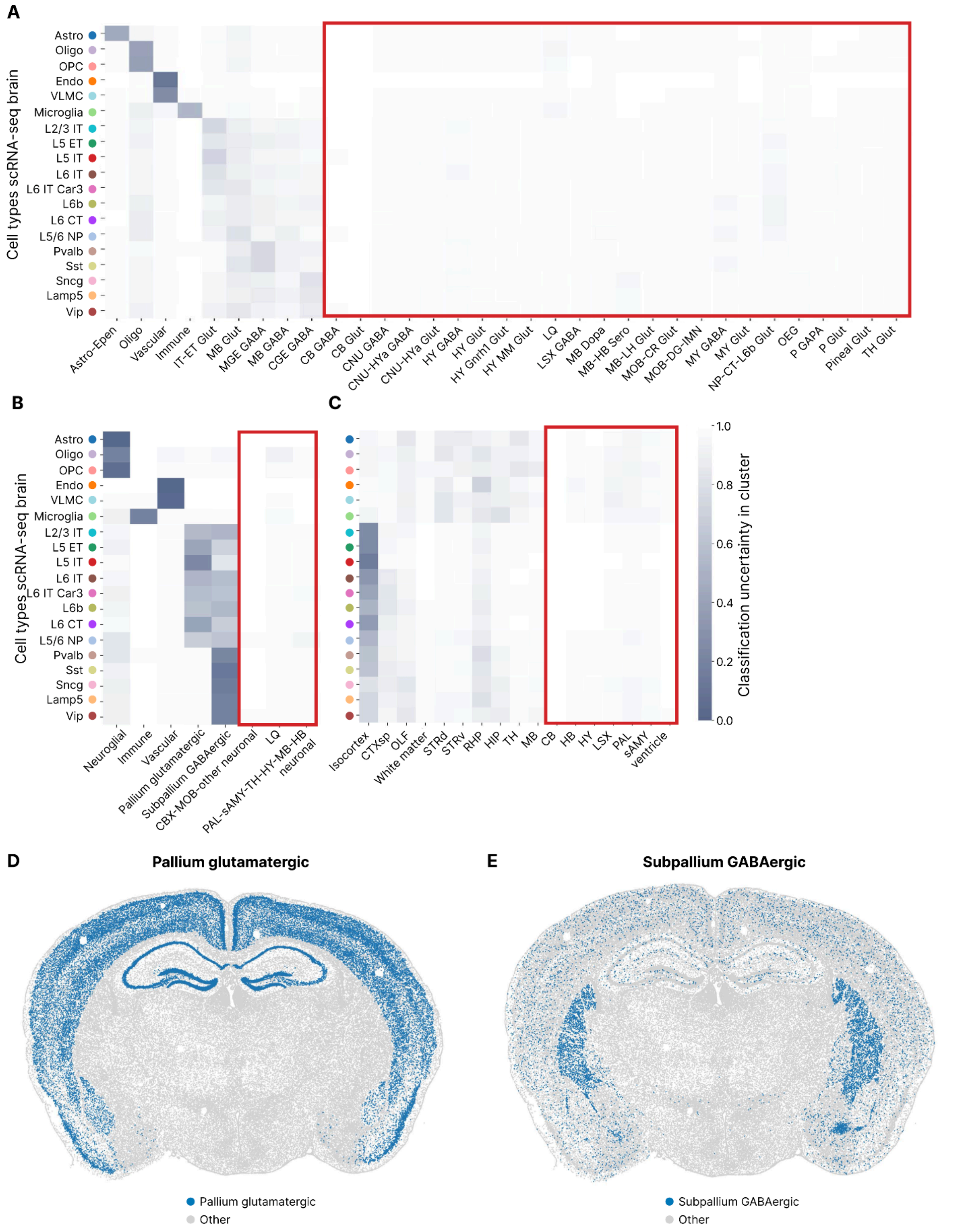


**Suppl. Figure 10 | Nicheformer label transfer classification uncertainty from spatial to dissociated assays. A-C)** Cell type (A), niche (B), and region (C) predicted label uncertainty across all cell types in the scRNA-seq mouse brain data. Nicheformer assigns lower uncertainty to plausible labels given the nature of the dataset and high uncertainty to labels not present in the primary motor cortex. The highlighted boxes show cell types, niches and regions one would not expect to find in the primary motor cortex. Nicheformer correctly shows a high uncertainty in those. D-E) Spatial allocation of cells in an exemplary section of the MERFISH mouse brain dataset colored by the pallium glutamatergic niche label (D) and the subpallium GABAergic niche label (E), respectively.

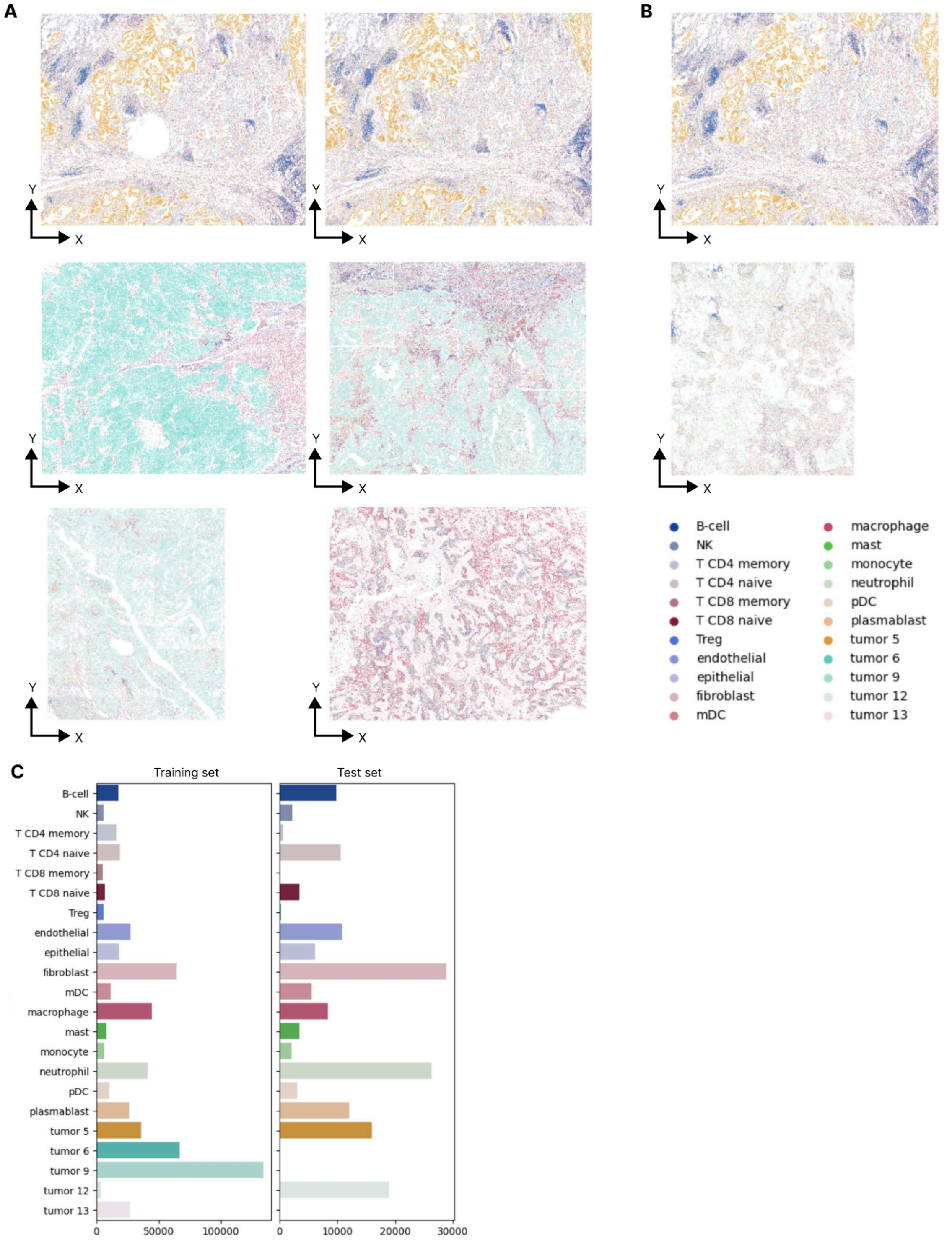


**Suppl. Figure 11 | Nicheformer fine-tuning datasets - CosMx human lung.** A-B) Spatial allocation of cells in the training set (A) and test set (B) tissue sections colored by cell type. C) Distribution of cell type labels in the training and test set in the CosMx human lung dataset.

### suSupplementary Data

| Hyperparameter | Value |
|---|---|
| Optimizer | [**AdamW**, Adam] |
| Optimizer momentum | $\beta_1 = 0.9$,<br>$\beta_2 = 0.99$ |
| Maximum learning rate | **1e-3** |
| Minimum learning rate | [**1e-5**, 0] |
| Weight decay | [**0.1**, 0.01, 0.0] |
| Dropout | [0.15, 0.10, **0.0**]] |
| Maximum gradient clipping | [**1.0**, 0.7, 0.5, 0.0] |
| Minimum gradient clipping | [**0.5**, 0.0] |
| Warmup iterations | **100K** |
| Batch size | **9** |
| Gradient accumulation | [**10**, 0] |
| Precision | [**bfloat16**, float32] |
| Embedding dimensionality | [**512**, 256] |
| FFN dimensionality | [**1024**, 512] |
| Transformer blocks | [**12**, 8, 6] |
| Attention heads | [**16**, 12, 8] |
| Gene tokens | **20310** |
| Contextual tokens | **15** |
| Number of parameters | [**49.3**, 40.9, 15.1] M |

**Suppl. Table 1 | Overview of pretraining architecture hyperparameters used for the final Nicheformer pretraining model.** Shown are the model hyperparameters screened for the Nicheformer pretraining with parameters highlighted in bold being the final parameter selection.

| | Nicheformer | Geneformer | CellPLM |
|---|---|---|---|
| **Dataset size** | SpatialCorpus-110M (110 million cells) | Genecorpus (30 million cells) | 11 million cells |
| **Tissue, species** | Cross-tissue, humans & mouse | Cross-tissue, human | Cross-tissue, human |
| **Modalities** | Dissociated & Spatial | Dissociated | Dissociated & Spatial |
| **Tokenization** | Gene rank-based value encoding | Gene rank-based value encoding | Cell tokenization |
| **Vocabulary size** | 20,310 protein-coding genes, (16,981 orthologous genes, 3,178 human-specific and 151 mouse-specific) | 25,424 protein-coding genes | 13, 500 common gene set |
| **Context length** | 1,500 | 2,048 | Uses a batch of cells as context |
| **Transformer units** | 12 | 6 | 4 |
| **Embedding dimension** | 512 | 256 | 512 |
| **Total parameters** | ~ 50 million | ~ 10 million | ~ 82 million |
| **Dissociated downstream tasks** | Cell type annotation | Gene function prediction, cell type annotation, GRN inference | Cell type annotation, genetic perturbation effect prediction |
| **Spatial downstream tasks** | Niche and region label prediction, neighborhood composition prediction, cellular density prediction | Not considered | Gene expression imputation |

**Suppl. Table 2 | Comparison of Nicheformer architecture to Geneformer and CellPLM.** Shown are the pretraining dataset specifications, model specifications and evaluated downstream tasks for Nicheformer compared to Geneformer and CellPLM.

| GEO ID | DOI | Number of cells | Tissue | Assay | Author |
|---|---|---|---|---|---|
| **GSE117824_human** | 10.1038/s41586-019-1367-0 | 51937 | blood; bone marrow | 10x 3' v2 | Nam_2019 |
| **GSE117824_mouse** | 10.1038/s41586-019-1367-0 | 738 | hematopoietic cell | 10x 3' v2 | Nam_2019 |
| **GSE118068** | 10.1038/s41586-019-1158-7 | 62893 | brain | 10x 3' v2 | Vladoiu_2019 |
| **GSE119940** | 10.1038/s41590-019-0403-4 | 46856 | spleen; bone marrow | 10x 3' v2 | Yao_2019 |
| **GSE124952** | 10.1038/s41467-019-12054-3 | 43795 | brain | 10x transcription profiling | Bhattacherjee_2019 |
| **GSE126060** | 10.1038/s41467-019-14172-4 | 41701 | tendon | 10x 3' v2 | Sorkin_2020 |
| **GSE128423** | 10.1016/j.cell.2019.04.040 | 135004 | bone marrow; bone tissue | 10x 3' v2 | Baryawno_2019 |
| **GSE128761** | 10.1016/j.stem.2020.08.001 | 231485 | liver; bone marrow | 10x 3' v2 | Li_2020 |
| **GSE128987** | 10.1038/s41556-020-00619-0 | 25932 | female reproductive system | 10x 3' v2 | Chumduri_2021 |
| **GSE129826** | 10.1016/j.celrep.2019.10.073 | 106670 | lymphoblast | 10x 3' v2 | Xie_2019 |
| **GSE130593** | 10.1242/dev.183251 | 53927 | testis | 10x 3' v2 | Tan_2020 |
| **GSE130822** | 10.1016/j.stem.2019.12.011 | 5401 | intestine | 10x 3' v3 | Murata_2020 |
| **GSE130879** | 10.1016/j.immuni.2019.12.002 | 31614 | lymph node; blood; adipose tissue; lung; skin of body; spleen; liver; bone marrow | 10x 3' v2 | Delacher_2020 |
| **GSE130888** | 10.1126/scitranslmed.aav5341 | 96446 | peritoneum | 10x 3' transcription profiling | Si_2019 |
| **GSE131339** | 10.1038/s41467-019-13465-y | 42213 | thymus | 10x 3' v2 | Cowan_2019 |
| **GSE131996** | 10.1016/j.immuni.2019.06.009 | 32805 | lung | 10x 3' v2 | Nagashima_2019 |
| **GSE132355** | 10.1038/s41467-020-18231-z | 341322 | hypothalamus | 10x transcription profiling | Kim_2020 |
| **GSE133531** | 10.1038/s41588-019-0531-7 | 65589 | brain | 10x 3' v2 | Jessa_2019 |
| **GSE134571** | 10.1038/s41586-019-1535-2 | 13661 | embryonic stem cell; amniotic stem cell | 10x 3' v2 | Zheng_2019 |
| **GSE135310** | 10.1161/CIRCRESAHA.120.317200 | 22630 | heart | 10x 3' v2; 10x 3' v3 | Vafadarnejad_2020 |
| **GSE135326** | 10.1038/s41586-0 | 280410 | brain | 10x 3' v3 | Chu_2019 |

| | | | | | |
|---|---|---|---|---|---|
| | 19-1644-y | | | | |
| **GSE135356** | 10.1038/s41467-020-18957-w | 60841 | endothelial cell of umbilical vein | 10x 3' v2 | Calandrelli_2020 |
| **GSE135414** | 10.1016/j.celrep.2020.01.075 | 10509 | retina | 10x 3' v2 | Jorstad_2020 |
| **GSE136394** | 10.1158/2326-6066.CIR-19-0299 | 73585 | intestine; blood | 10x 5' transcription profiling | Lu_2019 |
| **GSE136441** | 10.1073/pnas.2005570117 | 52542 | gonad; ovary | 10x 3' v2 | Niu_2020 |
| **GSE137026** | 10.1084/jem.20220126 | 43363 | lung | 10x 3' v2 | Liu_2022 |
| **GSE139168** | 10.1126/sciadv.aba9950 | 28731 | heart | 10x 3' v3 | Hatzistergos_2020 |
| **GSE140510** | 10.1038/s41591-019-0695-9 | 90647 | brain | 10x 5' transcription profiling | Zhou_2020 |
| **GSE140628** | 10.1158/2159-8290.CD-19-0958 | 14563 | pancreas | 10x transcription profiling | Zhang_2020 |
| **GSE141471** | 10.1038/s41467-021-27899-w | 54855 | hindlimb | 10x 3' v3 | Dutrow_2022 |
| **GSE141526** | 10.1126/sciadv.abm7981 | 56162 | skin of body | 10x 3' v2 | Yao_2020 |
| **GSE141552** | 10.1093/hmg/ddaa038 | 53616 | brain | 10x 3' v2 | Brenner_2020 |
| **GSE141784** | 10.1084/jem.20192362 | 106681 | pancreas | 10x 3' v2 | Zakharov_2020 |
| **GSE142143** | 10.1038/s41419-022-04693-0 | 30659 | brain | 10x 3' v2 | Lin_2022 |
| **GSE142797** | 10.1101/2020.04.27.063503 | 137873 | brain | 10x 3' v3 | Winkel_2020 |
| **GSE143293** | 10.1038/s41586-020-3017-y | 19345 | bone marrow | 10x transcription profiling | Yusufova_2020 |
| **GSE145216** | 10.1016/j.cell.2020.03.004 | 98191 | vagus nerve | 10x 3' v3 | Prescott_2020 |
| **GSE145251** | 10.1016/j.stem.2022.03.001 | 91242 | motor neuron; endodermal cell; cardiac muscle cell | 10x 3' v3 | Kong_2022 |
| **GSE145326** | 10.1172/JCI130323 | 63888 | breast | 10x 3' v3 | Garcia-Recio_2020 |
| **GSE145689** | 10.1681/ASN.2020070930 | 324635 | kidney | 10x 3' v2 | Hinze_2021 |
| **GSE145866** | 10.1016/j.celrep.2020.107952 | 420146 | intestine | 10x 3' v2 | Sheng_2020 |
| **GSE146122** | 10.1016/j.cell.2020.03.015 | 22194 | skin epidermis | 10x 3' v2 | Dekoninck_2020 |
| **GSE146138** | 10.1053/j.gastro.2020.09.011 | 110107 | intestine | 10x 3' v2 | Lähde_2021 |
| **GSE146194** | 10.1016/j.cell.2022.05.013 | 669018 | blood | 10x 3' v2 | Replogle_2020 |

| | | | | | |
|---|---|---|---|---|---|
| **GSE146298** | 10.1016/j.celrep.2020.03.059 | 159906 | brain | 10x 3' v2 | Darbandi_2020 |
| **GSE146512** | 10.1016/j.celrep.2020.108027 | 100639 | ovarian follicle | 10x 3' v3 | Man_2020 |
| **GSE148339** | 10.1016/j.jcmgh.2020.07.012 | 16714 | liver | 10x 3' v3 | Nault_2021 |
| **GSE148978** | 10.1016/j.immuni.2020.10.024 | 135297 | thymus | 10x 5' transcription profiling | Chopp_2020 |
| **GSE149040** | 10.1038/s41467-021-21704-4 | 86719 | retina | 10x 3' v2 | Wu_F_2021 |
| **GSE149201** | 10.1158/2159-8290.CD-20-0461 | 90263 | bone tissue | 10x 3' v3 | Khazaei_2020 |
| **GSE149356** | 10.1126/sciimmunol.abf0125 | 28775 | blood | 10x 5' transcription profiling | Tan_2021 |
| **GSE149931** | 10.1038/s41587-020-00763-w | 55202 | brain | 10x 3' v3 | Miura_2020 |
| **GSE150708** | 10.21203/rs.3.rs-62758/v1 | 32701 | lung | 10x 3' v2 | Duan_2020 |
| **GSE150871** | 10.1038/s41586-020-2795-6 | 127783 | spinal cord | 10x 3' v3 | Li_Y_2020 |
| **GSE150995** | 10.1172/JCI136142 | 13983 | tendon | 10x 3' v3 | Huber_2020 |
| **GSE151186** | 10.1016/j.stem.2020.10.003 | 5256 | embryonic stem cell | 10x 3' v3 | Mikryukov_2021 |
| **GSE152325** | 10.1038/s41556-020-00617-2 | 297808 | intestine | 10x 3' v2 | Böttcher_2021 |
| **GSE152573** | 10.1182/blood.2020007747 | 27176 | bone marrow | 10x 3' v2 | Zheng_2020 |
| **GSE152988** | 10.1038/s41593-021-00862-0 | 96637 | neuron | 10x 3' v3 | Tian_2021 |
| **GSE152999** | 10.1038/s41467-020-20351-5 | 174462 | intestine | 10x transcription profiling | Sarvestani_2021 |
| **GSE153099** | 10.1016/j.stemcr.2020.12.018 | 43143 | retina | 10x 3' v2 | Kruczek_2021 |
| **GSE153117** | 10.1038/s41467-021-26069-2 | 24057 | thymus | 10x 3' v2 | Cordero_2021 |
| **GSE153274** | 10.1038/s41467-021-22021-6 | 42996 | spermatogonium | 10x 3' v3 | Zhang_2021 |
| **GSE153288** | 10.1038/s41467-020-17544-3 | 39848 | thymus | 10x 3' transcription profiling | Lebel_2020 |
| **GSE153762** | 10.7554/eLife.60223 | 17384 | sciatic nerve | 10x 3' v3 | Kalinski_2020 |
| **GSE153770** | 10.1016/j.celrep.2020.108004 | 32412 | embryo; liver | 10x 3' v2; 10x 3' v3 | Simic_2020 |
| **GSE153802** | | 254501 | intestine | 10x 3' v3 | Hong_2020 |
| **GSE154196** | 10.15252/embj.2020106423 | 14296 | brain | 10x 3' transcription profiling | Jönsson_2021 |

| | | | | | |
|---|---|---|---|---|---|
| **GSE154359** | 10.1016/j.stem.2020.08.015 | 24845 | skin fibroblast | 10x 3' v2 | Cates_2021 |
| **GSE154386** | 10.1371/journal.ppat.1009240 | 200192 | blood | 10x 5' v2 | Waickman_2021 |
| **GSE154567** | 10.1016/j.celrep.2020.108590 | 85064 | blood | 10x 3' v3 | Yao_2021 |
| **GSE154579** | 10.1038/s41467-020-19234-6 | 97363 | skin of body | 10x 3' v2; 10x 3' transcription profiling | Lin_2020 |
| **GSE154932** | 10.1088/1478-3975/abb09c | 14964 | mammary gland epithelial cell | 10x 3' v2 | Johnson_2020 |
| **GSE155226** | 10.1126/scitranslmed.abf7872 | 47066 | heart | 10x 3' v3 | Pérez-Bermejo_2020 |
| **GSE155340** | 10.1126/sciimmunol.abb5168 | 5167 | bone marrow | 10x transcription profiling | Aykut_2020 |
| **GSE155788** | 10.7554/eLife.61413 | 35453 | brain | 10x 3' transcription profiling | Orsenigo_2020 |
| **GSE155850** | 10.1172/jci.insight.139932 | 88559 | skin of body | 10x 3' v3 | Lowe_2020 |
| **GSE156136** | 10.1016/j.cell.2020.09.062 | 38925 | bone marrow | 10x 3' transcription profiling | Khan_2020 |
| **GSE156183** | 10.1073/pnas.2017742118 | 94488 | brain | 10x 3' v2 | Ellwanger_2021 |
| **GSE156245** | 10.1038/s41586-021-03283-y | 23834 | colon | 10x 3' v2 | Wu_2021 |
| **GSE156285** | 10.1126/sciimmunol.abc6259 | 36091 | upper respiratory conduit | 10x 3' transcription profiling | Hung_2020 |
| **GSE156920** | 10.1038/s41467-021-23320-8 | 12967 | embryonic stem cell | 10x 3' v2 | Liu_2021 |
| **GSE157244** | 10.1161/JAHA.120.019019 | 153337 | blood; heart; bone marrow | 10x transcription profiling | Calcagno_2021 |
| **GSE157292** | 10.1172/jci.insight.141321 | 60181 | kidney | 10x transcription profiling | Dangi_2020 |
| **GSE157362** | 10.1242/dev.197111 | 3141 | testis | 10x 3' v3 | Webster_2021 |
| **GSE157525** | 10.1101/gad.339978.120 | 37665 | cerebral cortex | 10x 3' v2 | Parisian_2020 |
| **GSE157771** | 10.1038/s41593-020-00745-w | 102376 | brain; spleen | 10x 3' v2 | Jin_2021 |
| **GSE157773** | | 48705 | mesenchymal stem cell; embryonic stem cell; skeletal muscle satellite myogenic cell | 10x 3' transcription profiling | Han_2020 |
| **GSE157977** | 10.1126/science.aaz6063 | 89737 | brain | 10x 3' v2 | Jin_2020 |
| **GSE158038** | 10.1038/s41467-021-22210-3 | 96600 | blood; lung | 10x 5' transcription profiling | Ferreira-Gomes_2021 |

| | | | | | |
|---|---|---|---|---|---|
| **GSE158192** | 10.1038/s41467-021-22817-6 | 88389 | lung | 10x 3' v2 | Little_2021 |
| **GSE158356_mouse** | 10.26508/lsa.202000935 | 98297 | pancreas | 10x transcription profiling | Kemp_2021 |
| **GSE158450** | 10.1038/s41467-020-20343-5 | 29375 | brain | 10x 3' v2 | Joglekar_2021 |
| **GSE159354** | 10.1016/j.xcrm.2020.100140 | 184019 | lung | 10x 3' v2 | Gao_2020 |
| **GSE159519** | 10.1016/j.cell.2020.10.030 | 71279 | epithelial cell of lung | 10x 5' v1 | Daniloski_2021 |
| **GSE159977** | 10.1038/s41586-021-03362-0 | 112566 | liver | 10x 3' transcription profiling | Pfister_2021 |
| **GSE160061** | 10.1111/cpr.12933 | 11271 | germ cell | 10x 3' v3 | Yang_2021 |
| **GSE160097** | 10.1002/eji.202048797 | 35164 | blood; synovial fluid | 10x 5' transcription profiling | Maschmeyer_2021 |
| **GSE160098** | 10.1038/s41467-023-38647-7 | 68404 | myoblast | 10x 3' v3 | Sunadome_2023 |
| **GSE160664** | 10.1164/rccm.202008-3198OC | 7368 | lung | 10x 3' v2 | Okuda_2021 |
| **GSE160729** | 10.1016/j.cmet.2020.12.004 | 442800 | adipose tissue | 10x 3' v2 | Sárvári_2021 |
| **GSE160772** | 10.1096/fj.202002123R | 6379 | endometrium | 10x 3' v2 | Kirkwood_2021 |
| **GSE161066** | 10.3389/fphys.2021.637924 | 27111 | immature Schwann cell; amnion mesenchymal stem cell | 10x 3' v2 | Wei_2021 |
| **GSE161227** | 10.1084/jem.20212479 | 257991 | brain | 10x 3' v3 | Zhao_2022 |
| **GSE161230** | | 124799 | brain | 10x 3' v3 | Zhao_2020 |
| **GSE161363** | 10.1126/science.abc1944 | 85227 | epithelial cell of lung | 10x 3' transcription profiling | Quinn_2021 |
| **GSE161685** | 10.1172/jci.insight.144294 | 20071 | lung | 10x 3' transcription profiling | Gally_2021 |
| **GSE161937** | 10.1073/pnas.1915389116 | 41512 | uterus | 10x 3' v3 | Fitzgerald_2019 |
| **GSE162073** | 10.1084/jem.20200844 | 82138 | breast | 10x 3' v3 | Xu_2021 |
| **GSE162807_human** | 10.1038/s41467-022-28473-8 | 4605 | spinal cord | 10x 5' transcription profiling | Tansley_2022 |
| **GSE162807_mouse** | 10.1038/s41467-022-28473-8 | 239635 | spinal cord | 10x 5' transcription profiling | Tansley_2022 |
| **GSE163018** | 10.1038/s41421-021-00266-1 | 49534 | brain | 10x 3' transcription profiling | Ziffra_2021 |

| | | | | | |
|---|---|---|---|---|---|
| **GSE163278** | 10.1172/jci.insight.127807 | 63765 | bone marrow | 10x transcription profiling | Bailur_2019 |
| **GSE163650** | 10.1371/journal.pone.0244743 | 39794 | liver | 10x 3' v2 | Taylor_2021 |
| **GSE163668** | 10.1038/s41586-021-03234-7 | 189079 | blood | 10x 5' v1 | Combes_2021 |
| **GSE163701** | 10.1038/s41698-021-00160-9 | 9128 | adipose tissue | 10x 3' v3 | Su_2021 |
| **GSE163830** | | 49527 | adipose tissue | 10x 3' v3 | Gupta_2020 |
| **GSE163919** | | 12685 | lung | 10x 5' v1 | Banovich_2020 |
| **GSE164044** | 10.1016/j.neuron.2019.08.002 | 22559 | retina | 10x 3' v3 | Norrie_2019 |
| **GSE164573** | 10.1038/s41467-021-22842-5 | 34439 | skeletal muscle tissue | 10x 3' v2 | Julien_2021 |
| **GSE165551** | 10.7554/eLife.67436 | 54367 | brain | 10x 3' v2 | Cebrian-Silla_2021 |
| **GSE165554** | 10.7554/eLife.67436 | 35025 | brain | 10x 3' v3 | Cebrian-Silla_2021 |
| **GSE166218** | 10.1093/neuonc/noac138 | 161021 | brain | 10x 3' v2 | Friedrich_2023 |
| **GSE166262** | 10.1038/s41588-021-00818-x | 53214 | embryo | 10x 3' v3 | Kameneva_2021 |
| **GSE166525** | 10.1186/s13046-023-02686-1 | 113518 | brain | 10x 3' v3 | Liu_2023 |
| **GSE166797** | 10.1073/pnas.2023070118 | 7227 | lymphoblast | 10x transcription profiling | Wu_2021_2 |
| **GSE166992** | 10.1016/j.celrep.2021.108863 | 63895 | blood | 10x 5' transcription profiling | Thompson_2021 |
| **GSE167595** | 10.1158/1940-6207.CAPR-21-0378 | 62644 | intestine | 10x 5' transcription profiling | Yang_2022 |
| **GSE167992** | 10.1016/j.stem.2021.04.003 | 4787 | cornea | 10x 3' v2 | Altshulerls_2021 |
| **GSE168732** | 10.1038/s41467-021-25771-5 | 106453 | blood | 10x 3' transcription profiling | Wang_2021 |
| **GSE168758** | 10.1016/j.jhep.2021.03.029 | 101050 | liver | 10x 3' v3 | Reich_2021 |
| **GSE169718** | 10.1016/j.devcel.2021.12.012 | 51289 | intestine | 10x 3' v3 | Ohara_2022 |
| **GSE172127** | 10.1038/s41421-021-00266-1 | 18061 | liver | 10x 3' v2 | Lu_2021 |
| **GSE200218** | 10.1016/j.cell.2022.06.007 | 191032 | brain; axillary lymph node; subcutaneous adipose tissue | 10x 5' transcription profiling | Biermann_2022 |
| **GSE225278** | 10.1038/s41467-023-38704-1 | 201525 | pluripotent stem cell | 10x 3' v3 | Neavin_2023 |
| **GSE114687** | 10.1038/s41588-019-0489-5 | 75735 | cell in vitro | 10x 3' v1 | McFaline-Figueroa_2019 |

| | | | | | |
|---|---|---|---|---|---|
| **GSE117176** | 10.1172/jci.insight.126453 | 21574 | adipose tissue, bone marrow | 10x 3' v2 | Li_2019 |
| **GSE117770** | 10.1016/j.cmet.2019.01.021 | 30000 | pancreas | 10x 3' v2 | Thompson_2019 |
| **GSE120508** | 10.1038/s41422-018-0099-2 | 6199 | testis | 10x 3' v2 | Guo_2018 |
| **GSE122342** | 10.1016/j.stem.2018.12.015 | 11277 | dorsal plus ventral thalamus | 10x 3' v1 | Xiang_2019 |
| **GSE122960** | 10.1164/rccm.201712-2410OC | 80919 | lung | 10x 3' v2 | Reyfman_2018 |
| **GSE123722** | 10.1016/j.cell.2020.11.017 | 9770 | hindbrain | 10x 3' v3 | Andersen_2020 |
| **GSE124691** | 10.1016/j.celrep.2019.10.131 | 9804 | lymphoid system | 10x 3' v2 | Magen_2019 |
| **GSE128855** | 10.1038/s41593-019-0393-4 | 21966 | dura mater, brain, brain meninx, choroid plexus | 10x 3' v2 | Van Hove_2019 |
| **GSE129519** | 10.1038/s41586-019-1289-x | 166242 | telencephalon | 10x 3' v2 | Velasco_2019 |
| **GSE130238** | 10.1016/j.stem.2019.08.002 | 16086 | cerebral cortex | 10x 3' v2 | Trujillo_2019 |
| **GSE131685** | 10.1038/s41597-019-0351-8 | 25404 | kidney | 10x 3' v2 | Liao_2020 |
| **GSE132672** | 10.1038/s41586-020-1962-0 | 381350 | prefrontal cortex, hippocampal formation, parietal cortex, occipital cortex, primary motor cortex, primary visual cortex, somatosensory cortex, frontal cortex, cerebral cortex, telencephalon, temporal cortex | 10x 3' v2 | Bhaduri_2020 |
| **GSE135893** | 10.1126/sciadv.aba1972 | 114396 | lung parenchyma | 10x 3' v2", "10x 5' v1 | Habermann_2020 |
| **GSE136001** | 10.1038/s41467-021-21407-w | 41059 | brain | 10x 3' v2", "10x 3' v3 | Ochoka_2021 |
| **GSE136103** | 10.1038/s41586-019-1631-3 | 58358 | liver | 10x 3' v2 | Ramachandran_2019 |
| **GSE136831** | 10.1126/sciadv.aba1983 | 312928 | lung | 10x 3' v2 | Adams_2020 |
| **GSE143317** | 10.1038/s41591-021-01245-5 | 8020 | bone marrow | 10x 3' v2 | Da Via_2021 |
| **GSE145122** | 10.1038/s41591-020-1043-9 | 45300 | cerebral cortex | 10x 3' v2 | Khan_2020 |
| **GSE147405** | 10.1038/s41467-020-16066-2 | 97971 | cell in vitro | 10x 3' v2", "10x 3' v3 | Cook_2020 |
| **GSE149931** | 10.1038/s41587-020-00763-w | 27601 | ventral part of telencephalon | 10x 3' v3 | Miura_2020 |

| | | | | | |
|---|---|---|---|---|---|
| **GSE150903** | 10.1126/science.aaz5626 | 32464 | choroid plexus, telencephalon | 10x 3' v3 | Pellegrini_2020 |
| **GSE156728** | 10.1126/science.abe6474 | 183913 | kidney, esophagus, skin epidermis, endometrium, ovary, fallopian tube, thyroid gland, blood, pancreas | 10x 5' transcription profiling | Zheng_2022 |
| **GSE156989** | 10.1016/j.isci.2021.102404 | 487890 | blood | 10x 3' v3 | Savage_2021 |
| **GSE160641** | 10.1126/sciadv.abg9518 | 10973 | mammary gland | 10x 3' v3 | Opzoomer_2021 |
| **GSE161125** | 10.1038/s41467-020-20540-2 | 6904 | bone marrow | 10x 3' transcription profiling | Munoz-Rojas_2021 |
| **GSE163973** | 10.1038/s41467-021-24110-y | 40655 | dermis | 10x 3' v3 | Deng_2021 |
| **GSE164690** | 10.1038/s41467-021-27619-4 | 151680 | neck | 10x 3' v2 | Kuerten_20221 |
| **GSE165577** | 10.1038/s41593-021-00906-5 | 58586 | brain | 10x 3' v3 | Samarasinghe_2021 |
| **GSE168323** | 10.1038/s41467-021-27464-5 | 123294 | midbrain tegmentum | 10x 3' v3 | Fiorenzano_2021 |
| **GSE173351** | 10.1038/s41586-021-03752-4 | 902463 | blood, lung, brain, lymph node | 10x 5' transcription profiling | Caushi_2021 |
| **GSE187877** | 10.1038/s41586-021-04330-4 | 6899 | telencephalon | 10x 3' v3 | Kelava_2022 |
| **GSE190604** | 10.1126/science.abj4008 | 103805 | blood | 10x 3' v3 | Schmidt_2022 |
| **GSE201349** | 10.1038/s41588-022-01088-x | 826250 | colon | 10x 3' v3 | Becker_2022 |
| **GSE205554** | 10.15252/embj.2022111118 | 12310 | cerebral cortex | 10x 3' v3 | Vertesy_2022 |
| | 10.1126/science.adf1226 | 1665937 | entorhinal cortex, head, midbrain, medulla oblongata, diencephalon, cortex, hypothalamus, telencephalon, forebrain, hippocampal formation, parietal cortex, occipital cortex,'dorsal plus ventral thalamus', cerebellum', striatum', 'pons', temporal cortex, frontal cortex, hindbrain, cerebral subcortex, | 10x 3' v2", "10x 3' v3 | Brau_2023 |

| | | | | | |
|---|---|---|---|---|---|
| | | | telencephalon neural crest, brain, caudate-putamen | | |
| | 10.1016/j.devcel.2020.01.033 | 36807 | lung | 10x 3' v2 | Miller_2020 |
| | 10.1038/s41586-022-05279-8 | 34088 | brain | 10x 3' v3 | Fleck_2022 |
| | 10.1016/j.cell.2022.09.010 | 348379 | cortical plate, brain telencephalon | 10x 3' v2", "10x 3' v3 | Uzquiano_2022 |
| | 10.1038/s41586-021-04358-6 | 663879 | telencephalon | 10x 3' v2", "10x 3' v3 | Paulsen_2022 |
| | 10.1158/2159-8290.CD-21-0316 | 178630 | liver | 10x 3' v3 | Wu_2022 |
| | 10.1186/s13059-020-02147-4 | 11512 | brain | 10x 3' v2 | He_2020 |
| | 10.1016/j.cell.2020.08.001 | 102203 | blood | 10x 3' v3 | Schulte-Schrepping_2020 |
| | 10.1371/journal.ppat.1008408 | 15085 | blood | 10x 3' v2 | de Vries_2020 |
| | 10.1038/s41467-020-15543-y | 43112 | blood | 10x 3' v2 | Cano-Gamez_2020 |
| | 10.1038/s41592-021-01344-8 | 87051 | brain | 10x 3' v3 | He_2022 |
| | 10.1038/s41586-019-1654-9 | 90576 | brain | 10x 3' v2 | Kanton_2019 |
| | 10.1016/j.cell.2021.11.033 | 54975 | lung | 10x 3' v3 | Wendisch_2021 |
| | 10.1038/s41587-021-01139-4 | 44429 | brain | 10x 3' v3 | Kleshchevnikov_2020 |
| | 10.1016/j.medj.2022.05.002 | 102441 | minor salivary gland, lung, synovial membrane of synovial joint, large intestine | 10x 3' v2", "10x 3' v3 | Korsunsky_2022 |

**Suppl. Table 3 | Overview of the additional dissociated data extracted and harmonized from GEO and sfaira.** Shown is the GEO ID used for downloading the data, the DOI, the number of cells before filtering low-quality cells, the primary tissue measured, the assay and the first author and publication year of the dataset and the respective publication.

| Author/ Name | DOI / Link | Number of cells | Tissue | Assay | Species | N sections / N donors |
|---|---|---|---|---|---|---|
| Garrido-Trigo et al. | 10.1038/s41467-023-40156-6 | 463,967 | Bowel | CosMx | Human | 171 / 9 |
| Nanostring CosMx mouse brain | https://nanostring.com/products/cosmx-spatial-molecular-imager/ffpe-dataset/ | 86,916 | Brain | CosMx | Mouse | 130 / 2 |
| Nanostring CosMx human liver | https://nanostring.com/products/cosmx-spatial-molecular-imager/ffpe-dataset/ | 793,318 | Liver | CosMx | Human | 2 / 2 |
| Nanostring CosMx human lung | https://nanostring.com/products/cosmx-spatial-molecular-imager/ffpe-dataset/ | 771,236 | Lung | CosMx | Human | 8 / 5 |
| SciLifeLab ISS mouse brain | Unpublished | 592,421 | Brain | ISS | Mouse | 20 / 13 |
| SciLifeLab ISS human brain | Unpublished | 988,901 | Brain | ISS | Human | 13 / 13 |
| SciLifeLab ISS human lung | Unpublished | 260,398 | Lung | ISS | Human | 14 /14 |
| Fang et al. | 10.1126/science.abm1741 | 127,273 | Brain | MERFISH | Human / Mouse | 378 / 6 |
| Allen et al. | 10.1016/j.cell.2022.12.010 | 378,918 | Brain | MERFISH | Mouse | 783 / 12 |
| Allen Institute brain atlas mouse p20 | Unpublished | 4,753,243 | Brain | MERFISH | Mouse | 59 / 1 |
| Allen Institute brain atlas mouse p28 | Unpublished | 4,064,846 | Brain | MERFISH | Mouse | 55 / 1 |
| Allen Institute brain atlas mouse female | Unpublished | 4,925,098 | Brain | MERFISH | Mouse | 54 / 1 |
| Vizgen MERFISH mouse brain | https://vizgen.com/data-release-program/ | 734,696 | Brain | MERFISH | Mouse | 9 / 3 |
| Androvic et al. | 10.1038/s41467-023-39447-9 | 347,968 | Brain | MERFISH | Mouse | 1646 / 3 |
| Zhang et al. | 10.1038/s41586-021-03705-x | 276,556 | Brain | MERFISH | Mouse | 5000 / 2 |
| Zhang et al. | 10.1038/s41586-023-06808-9 | 9,343,457 | Brain | MERFISH | Mouse | 2151 / 4 |
| Yao et al. | 10.1038/s41586-023-06812-z | 4,334,174 | Brain | MERFISH | Mouse | 59 / 1 |
| Vizgen MERFISH | ttps://vizgen.com/data-release-pro | 713.121 | Breast | MERFIS | Human | 1 / 1 |

| | | | | | | |
|---|---|---|---|---|---|---|
| human breast cancer | gram/ | | | H | | |
| Vizgen MERFISH human colon cancer | https://vizgen.com/data-release-program/ | 1,495,039 | Colon | MERFISH | Human | 2 / 2 |
| Vizgen MERFISH human liver cancer | https://vizgen.com/data-release-program/ | 1,166,496 | Liver | MERFISH | Human | 2 / 2 |
| Vizgen MERFISH human lung cancer | https://vizgen.com/data-release-program/ | 1,190,501 | Lung | MERFISH | Human | 2 / 2 |
| Vizgen MERFISH human ovarian cancer | https://vizgen.com/data-release-program/ | 896,638 | Ovary | MERFISH | Human | 4 / 2 |
| Vizgen MERFISH human prostate cancer | https://vizgen.com/data-release-program/ | 1,715,493 | Prostate | MERFISH | Human | 2 / 2 |
| Vizgen MERFISH human skin cancer | https://vizgen.com/data-release-program/ | 676,007 | Skin | MERFISH | Human | 2 / 2 |
| Vizgen MERFISH human uterine cancer | https://vizgen.com/data-release-program/ | 2,343,540 | Uterus | MERFISH | Human | 3 / 2 |
| Choi et al. | 10.1038/s41467-023-40674-3 | 367,369 | Retina | MERFISH | Mouse | 17 / 12 |
| Magen et al. | 10.1038/s41591-023-02345-0 | 1.671.375 | Liver | MERFISH | Human | 10 / 7 |
| 10x Genomics Xenium human brain | https://www.10xgenomics.com/datasets | 110248 | Brain | Xenium | Human | 3 / 3 |
| 10x Genomics Xenium mouse brain - Alzheimers | https://www.10xgenomics.com/datasets | 351.714 | Brain | Xenium | Mouse | 6 / 6 |
| 10x Genomics Xenium mouse brain | https://www.10xgenomics.com/datasets | 474.734 | Brain | Xenium | Mouse | 3 / 1 |
| 10x Genomics Xenium breast cancer (add-on) | https://www.10xgenomics.com/datasets | 2,721,056 | Breast | Xenium | Human | 4 / 3 |
| 10x Genomics Xenium colon cancer (add-on) | https://www.10xgenomics.com/datasets | 1,234,639 | Colon | Xenium | Human | 2 / 2 |
| 10x Genomics Xenium healthy colon (add-on) | https://www.10xgenomics.com/datasets | 546,806 | Colon | Xenium | Human | 2 / 2 |
| 10x Genomics Xenium kidney cancer (add-on) | https://www.10xgenomics.com/datasets | 56.510 | Kidney | Xenium | Human | 1 / 1 |

| | | | | | | |
|---|---|---|---|---|---|---|
| 10x Genomics Xenium healthy kidney (add-on) | https://www.10xgenomics.com/datasets | 97.560 | Kidney | Xenium | Human | 1 / 1 |
| 10x Genomics Xenium lung cancer (add-on) | https://www.10xgenomics.com/datasets | 681,530 | Lung | Xenium | Human | 2 / 2 |
| 10x Genomics Xenium healthy lung (add-on) | https://www.10xgenomics.com/datasets | 295.883 | Lung | Xenium | Human | 1 / 1 |
| 10x Genomics Xenium lymph node | https://www.10xgenomics.com/datasets | 377.985 | Lymph Node | Xenium | Human | 1 / 1 |
| 10x Genomics Xenium pancreas cancer | https://www.10xgenomics.com/datasets | 190.965 | Pancreas | Xenium | Human | 1 / 1 |
| 10x Genomics Xenium healthy pancreas | https://www.10xgenomics.com/datasets | 103901 | Pancreas | Xenium | Human | 1 / 1 |
| 10x Genomics Xenium skin cancer (add-on) | https://www.10xgenomics.com/datasets | 194,479 | Skin | Xenium | Human | 2 / 2 |
| SciLifeLab Xenium mouse brain | Unpublished | 624,058 | Brain | Xenium | Mouse | 1 / 1 |
| Kukanja et al. | 10.1016/j.cell.2024.02.030 | 660,801 | Spinal Cord | Xenium | Human | 6 / 6 |

**Suppl. Table 4 | Overview of the spatial datasets extracted and harmonized.** Shown is the author name of the original publication or the name describing the dataset, the DOI or dataset retrieval link used for downloading the data, the number of cells, tissue, assay, species, and the number of tissue sections and donors/ animals measured in the dataset. For the overview datasets obtained via the Vizgen data release and the 10x Genomics data resource are grouped by tissue and condition to reduce table size. Datasets highlighted in blue mark the dataset used for downstream Nicheformer tasks.

| Hyperparameter | Nicheformer (fine-tuned) & Nicheformer (linear probing) | Nicheformer (MLP) | scVI (linear probing) | PCA (linear probing) |
|---|---|---|---|---|
| Optimizer | AdamW | Adam | Adam | Adam |
| Optimizer momentum | $\beta_1 = 0.9$<br>$\beta_2 = 0.99$ | $\beta_1 = 0.9$<br>$\beta_2 = 0.99$ | $\beta_1 = 0.9$<br>$\beta_2 = 0.99$ | $\beta_1 = 0.9$<br>$\beta_2 = 0.99$ |
| Maximum learning rate | 1e-4 | 1e-3 | 1e-3 | 1e-3 |
| Minimum learning rate | 1e-5 | 1e-3 | 1e-3 | 1e-3 |
| Cosine decay length | 1 epoch | - | - | - |
| Weight decay | 0.1 | 0001 | 0.001 | 0.001 |
| Dropout | 0.0 | - | - | - |
| Maximum gradient clipping | 1.0 | - | - | - |
| Batch size | 9 | 256 | 256 | 256 |
| Gradient accumulation | 10 | - | - | - |
| Hidden dimensions | - | 256 | - | - |
| Precision | bfloat16 | 16-mixed | 16-mixed | 16-mixed |
| Epochs | 1 | 5 | 1 | 1 |

**Suppl. Table 5 | Overview of the fine-tuning and linear probing hyperparameters.** Shown are the model hyperparameters used for the different Nicheformer downstream tasks and the model parameters used for MLP and linear probing models trained on scVI and PCA embeddings.
